# A frameshift mutation is repaired through nonsense-mediated gene revising in *E. coli*

**DOI:** 10.1101/069971

**Authors:** Xiaolong Wang, Xuxiang Wang, Chunyan Li, Haibo Peng, Yalei Wang, Gang Chen, Jianye Zhang

## Abstract

The molecular mechanisms for repairing DNA damages and point mutations have been well understood but it remains unclear how a frameshift mutation is repaired. Here we report that frameshift reversion occurs in *E. coli* more frequently than expected and appears to be a targeted gene repair signaled by premature termination codons (PTCs), producing high-level variations in the repaired genes. Genome resequencing shows that the revertant genome is highly stable, and the single-molecule variations in the repaired genes are derived from RNA editing. A multi-omics analysis shows that the expression levels change greatly in most the DNA and RNA manipulating genes. DNA replication, transcription, RNA editing, RNA degradation, nucleotide excision repair, mismatch repair, and homologous recombination were upregulated in the frameshift or revertant, but the base excision repair was not. Moreover, genes and transposons in a duplicate region silenced in wild type *E. coli* were activated in the frameshift. Finally, we propose a *nonsense-mediated gene revising* (NMGR) model for frame repair, which also acts as a driving force for molecular evolution. In essence, nonsense mRNAs are recognized, edited, and transported to template the repair of the coding gene by RNA-directed DNA repair, nucleotide excision, mismatch repair, and homologous recombination. Thanks to NMGR, the mutation rate temporarily rises in a frameshift gene, bringing genetic diversity while repairing the frameshift mutation and accelerating the evolution process without a high mutation rate in the genome.

## 1. Introduction

DNA replication and DNA repair happen in every cell division and multiplication. Physical or chemical mutagens, such as radiation, pollutants, or toxins, can induce point mutations, insertions/deletions (InDels), and damages in DNA. Besides, because of the imperfect fidelity of DNA polymerase, spontaneous mutations and InDels also occur as replication errors or slipped-strand mispairing. If an InDel happens in a coding DNA sequence (CDS), and its size is not a multiple of three, it causes a frameshift mutation, leading to a substantial change in its encoded amino acid sequence. Besides, premature termination codons (PTCs) are often produced at the downstream of the InDel, resulting in truncated and dysfunctional proteins [1], leading to genetic disorders or even death.

The repair of DNA damages and point mutations have been intensively studied [2], including base-/nucleotide-excision, mismatch repair, homologous recombination, and non-homologous end-joining. However, it remains unclear how frameshift mutation is repaired. The reverse mutation phenomenon was discovered as early as in the 1960s [3], but it has long been explained simply by spontaneous mutagenesis (Fig 1). In principle, the reverse mutation rate is much lower than the forward mutation rate. However, it was reported that frameshift reversion occurs more frequently than expected in *Escherichia coli* (*E. coli*), which was believed to be an adaptive mutation [4-8]. Here we report that frameshift reversion occurs in *E. coli* much more frequently than expected and appears to be a targeted gene repair involving many gene repairing pathways.

**Fig 1.**
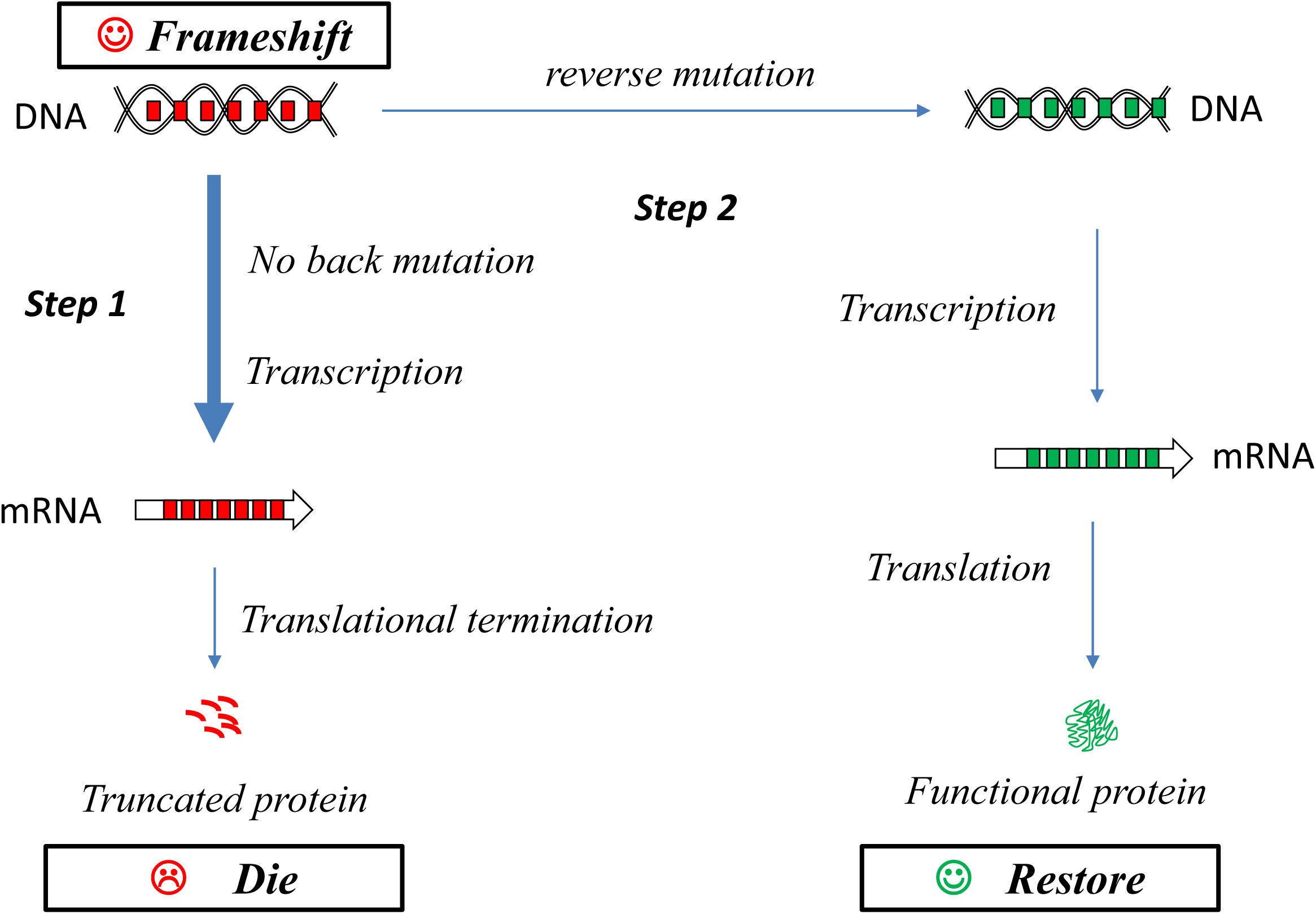
The traditional model for frameshift reversion. *Step 1*: when a frameshift mutation occurs in a CDS, many stop codons (*red bars*) emerge in the CDS, and PTCs in the nonsense mRNAs cause translational termination and truncated products; *Step 2*: the spontaneous mutagenesis causes a reverse mutation, repairs the frameshift gene, the reading frame restores, the PTCs hide (*green bars*), transcription and translation resume, producing functional mRNAs and proteins, the gene restores, and the cell recovers.

## 2. Materials and Methods

### 2.1 Frameshift preparation and revertant screening

The G:C base pair at +136 was deleted from the wild-type *bla* gene (*bla+*) by using an overlapping extension polymerase chain reaction (OE-PCR), resulting in a plasmid containing a frameshift *bla* gene, pBR322-(*bla*-). Competent cells of *E. coli* DH5α were transformed with plasmid pBR322 or pBR322-(*bla*-), propagated in tetracycline broth; dilutions were plated in parallel on tetracycline plates and ampicillin plates to screen for revertants. The recovery rates were calculated by a standard method [9]. The growth rates were evaluated by the doubling time in the exponential growth phase. The plasmid DNA was extracted, and the *bla* genes were sequenced by the Sanger method.

### 2.2 Construction and expression of a PTC-substituted frameshift gene

A PTC-substituted *bla-* gene, denoted as *bla*#, was derived from *bla-* by replacing each nonsense codon with a sense codon according to the readthrough rules (Table 1). A stop codon, TAA, was added at the 3’-end. The *bla*# gene was chemically synthesized by *Sangon Biotech, Co. Ltd* (*Shanghai*), inserted into the expression vector pET28a and transformed into *E. coli* competent cells strain BL21. The transformants were plated on a kanamycin plate, propagated in kanamycin broth, and plated on ampicillin plates to screen for revertants. The expression of the *bla#* gene was induced by 0.1 mM IPTG. The total protein samples were analyzed by sodium dodecyl sulfate-polyacrylamide gel electrophoresis (SDS-PAGE), and the product protein was purified by a nickel column chromatography. The purified product was tested by an iodometry assay to measure its lactamase activity [10].

**Table 1.** The readthrough rules derived from natural suppressor tRNA genes

| Gene | tRNA (AA) | Codon |
| --- | --- | --- |
| *supD* | Ser (**S**) | UAG |
| *supE* | Gln (**Q**) | UAG |
| *supF* | Tyr (**Y**) | UAG |
| *supG* | Lys (**K**) | UAA |
| *supU* | Trp (**W**) | UGA |

### 2.3 Genome resequencing and variation analysis

Genomic DNA was extracted from the frameshift (*Fs*) and the revertant (*Rs*). The *Novogene Co. Ltd.* performed next-generation sequencing (NGS) on the Illumina HiSeq 250PE platform. For each strain, clean reads were mapped onto the reference sequence of *E. coli* K12 MG1655 (NC_000913.3) and pBR322 (J01749.1). Single nucleotide polymorphisms (SNPs), InDels, and structural variations (SVs) were analyzed.

### 2.4 Transcriptome profiling and gene expression analysis

Total RNA samples were extracted from four different strains, including a wild type (*Wt*), a frameshift (*Fs*), a slow-growing revertant (*Rs*), and a fast-growing revertant (*Rf*). *Novogene Co. Ltd* performed RNA sequencing (RNAseq). The expected number of Fragments PerKilobase of transcript sequence per million base pairs sequenced (FPKM) was calculated for each gene. The threshold for significantly differential expression was set as corrected P-value (*q*) < 0.005 and fold change (*f*) ≥ 2.0. The enrichment analysis of Gene Ontology (GO) terms and Kyoto encyclopedia of genes and genomes (KEGG) pathways for the identified DEGs were performed.

### 2.5 Integrative analysis of genome and transcriptome datasets

All NGS datasets for wild-type *E. coli* that are available in Sequence Read Archive (SRA) were downloaded, including 1 genomic (SRR11474703) and 7 transcriptomic or RNAseq (SRR1149439, SRR2914524, SRR2914548, SRR2914549, SRR2914550, SRR6111094, and SRR6111095) datasets; the RNAseq datasets are pooled into a large transcriptomic dataset for wild-type *E. coli*. After mapping the reads onto the reference genome, the genomic and the transcriptomic coverage depths for each location (in KB) were plotted next to each other on a circular map using Circos (v0.69) [11].

### 2.6 Quantitative analysis of global proteomes

Total protein samples were prepared for the frameshift (*Fs*) and the revertant (*Rs*). Quantitative analysis of the proteome was performed using a service provided by *PTM-Biolabs* (*Hangzhou*), *Co., Ltd*. Identified protein IDs were converted into UniProt ID and mapped to GO IDs. For the identified proteins unannotated in the UniProt-GOA database, InterProScan is used to determine their GO annotations.

## 3. Results and Analysis

### 3.1 The growth of the frameshift and the revertants

The plasmid pBR322 contains two wild-type resistance genes, a β-lactamase gene (*bla+*) and a tetracycline resistance gene (*tet+*). The +136 G:C base pair of the *bla* gene was deleted by OE-PCR (Fig 2A), resulting in a plasmid containing a frameshift *bla*, pBR322-(*bla*-). This deletion is a lost-of-function mutation because 17 PTCs appeared, and all the active sites locates at the downstream of the deletion, including the acyl ester intermediate (AA 68), the proton receptor (AA 166), and the substrate-binding site (AA 232-234), which were all destroyed.

**Fig 2.**
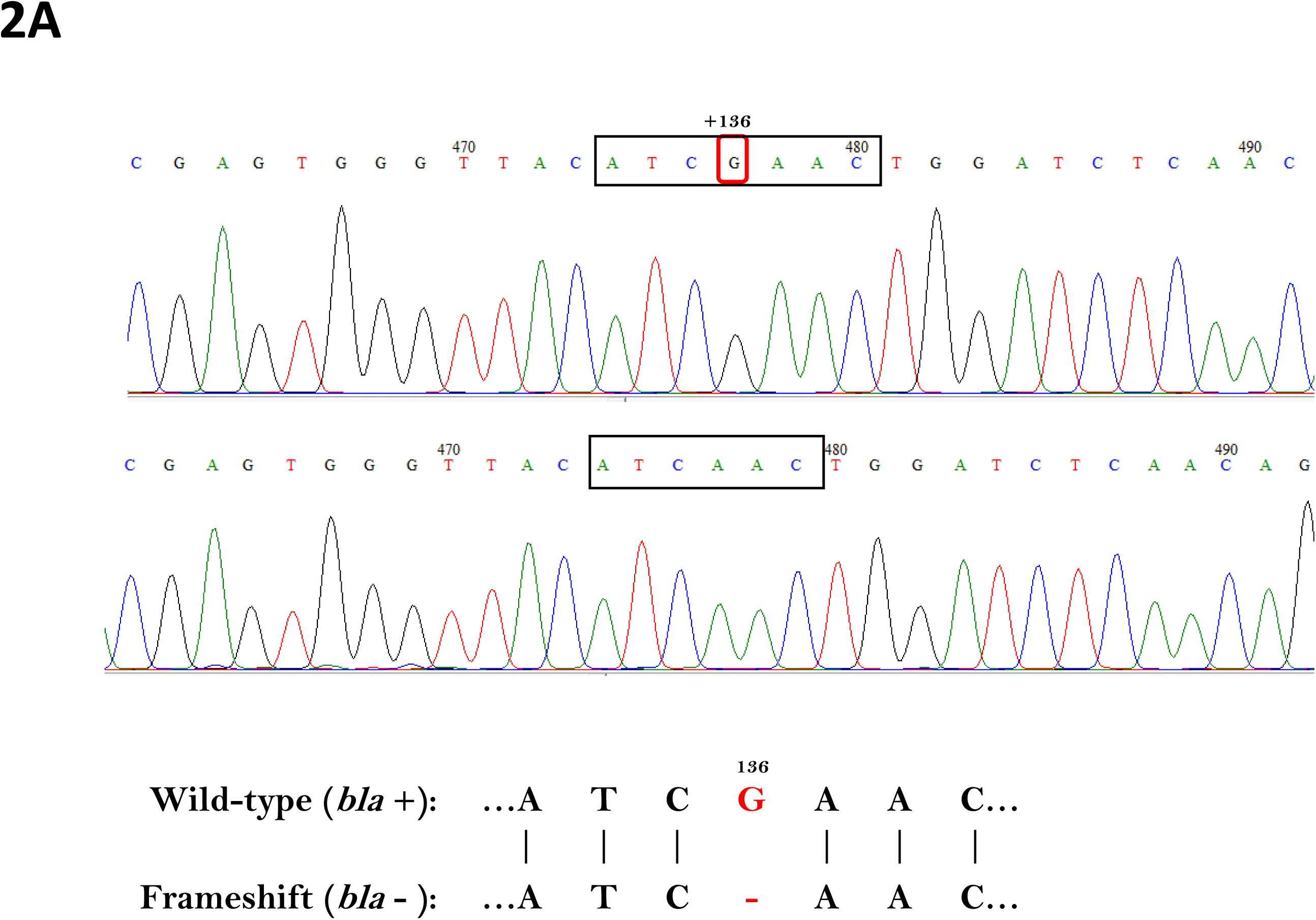
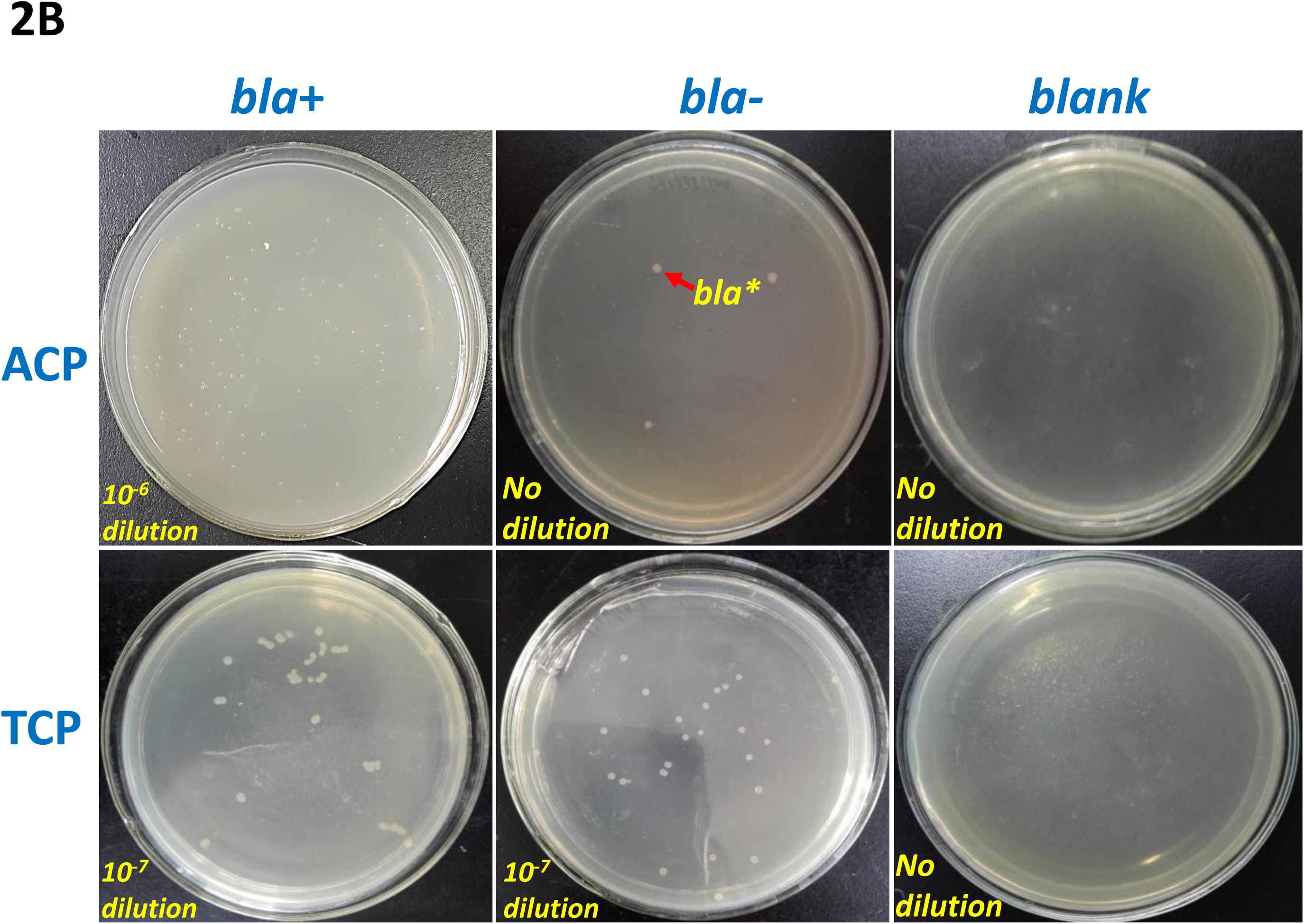
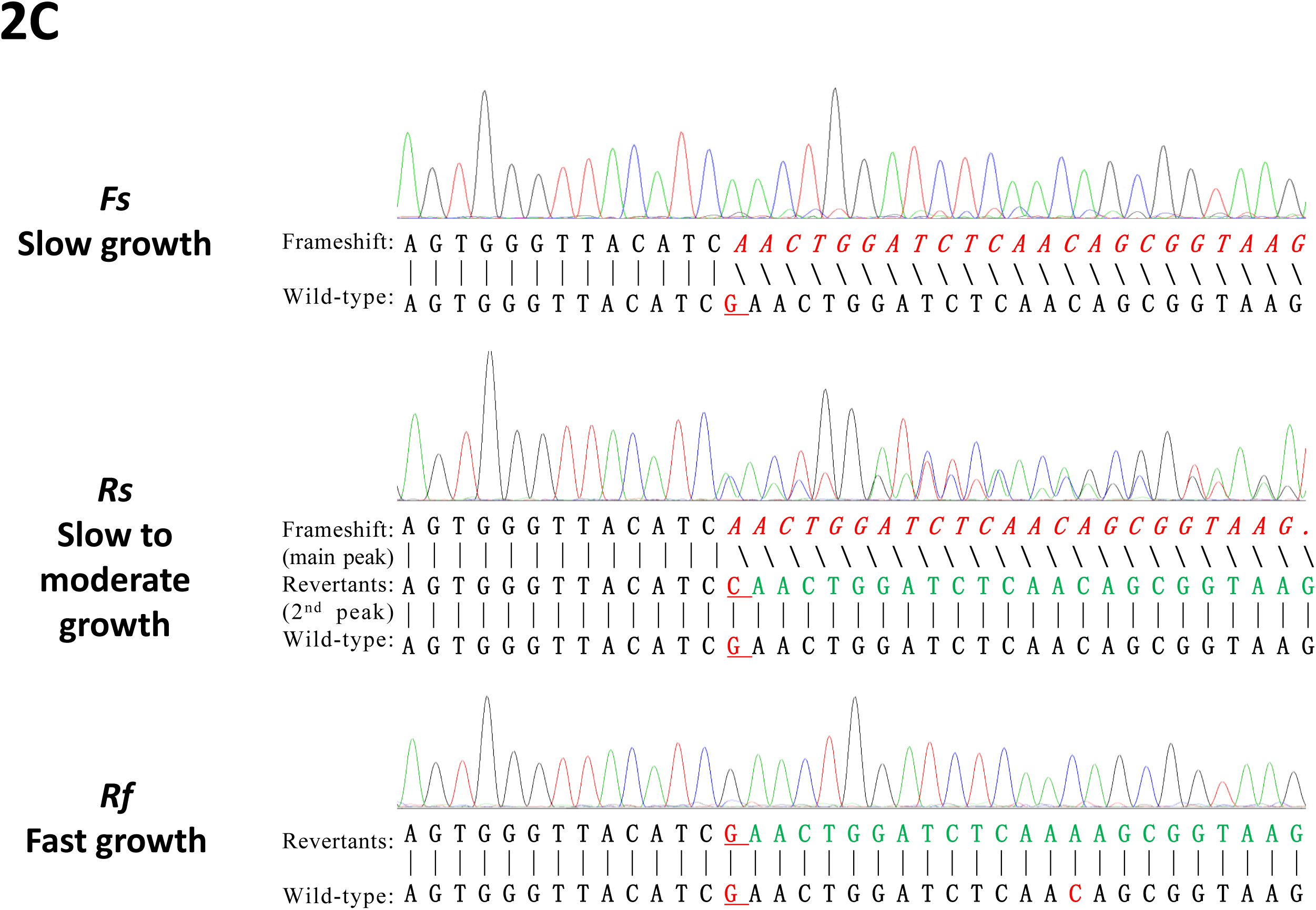
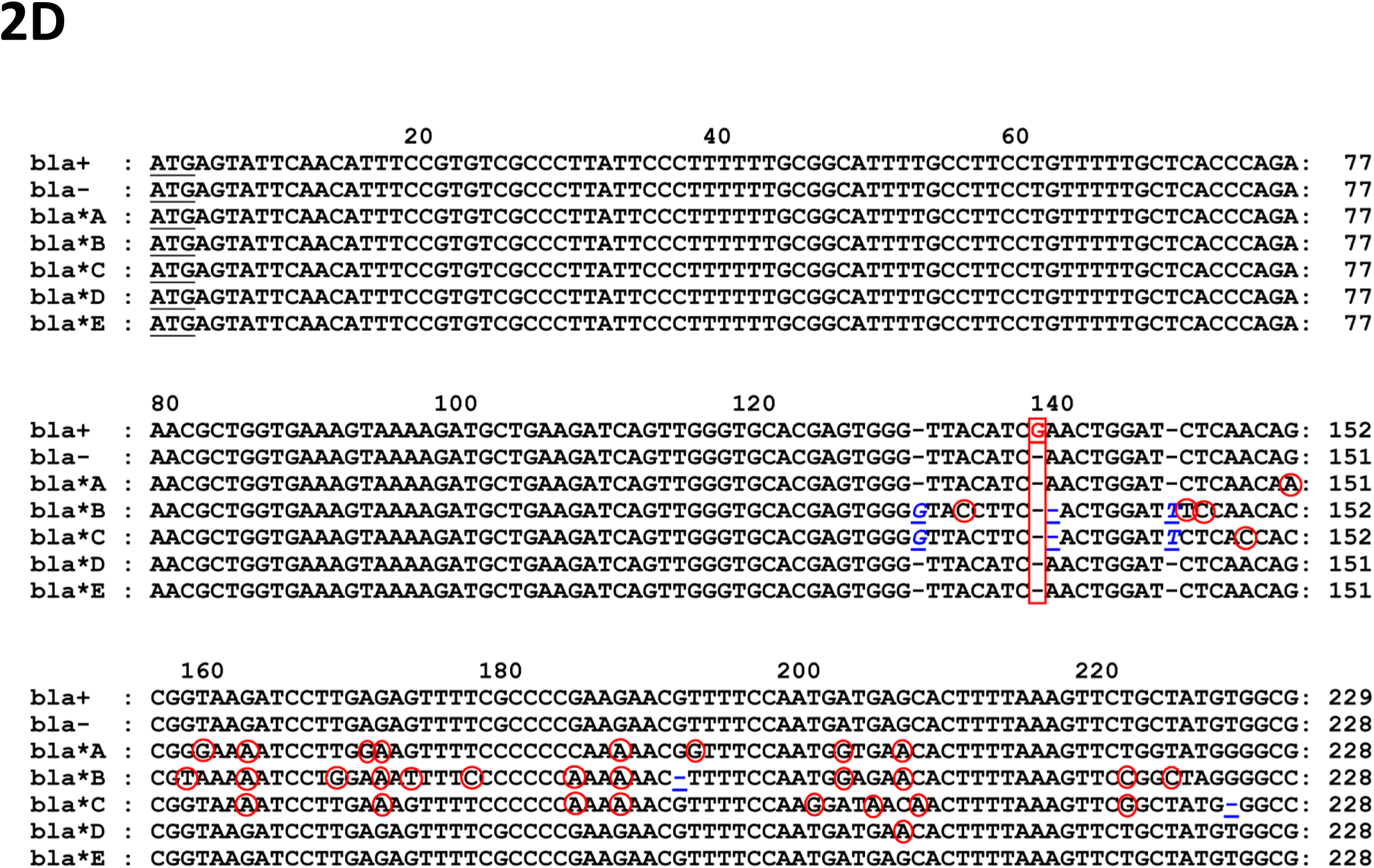
Growth of the wild type, the frameshift, and the revertants and Sanger sequencing of their *bla* genes: **(A)** The introduction of a frameshift mutation of the *bla* gene in the plasmid pBR322. *Top*: Sanger sequencing of the wild-type (*bla*+); *center*: Sanger sequencing of the frameshift (*bla-*); *bottom*: the alignment of *bla+* and *bla*-sequences. **(B)** Growth of the wild type, the frameshift, and the revertants on ampicillin- or tetracycline plates: *bla*+: wild type; *bla*-: frameshift; *bla*\*: revertants; *blank*: blank control; ACP: ampicillin plates; TCP: tetracycline plates. **(C)** The Sanger sequencing diagrams of the *revertants’ bla gene*s: *Top*: an initial revertant, the *bla* gene is still frameshifted. *Center*: a slow-growing revertant (*Rs*), its sequencing diagram contains two sets of overlapping peaks, the main and the secondary peaks represent the frameshift (*bla-*) and the repaired (*bla\**) gene, respectively. *Underlined:* showing a reacquired base C is inserted in the place of the deletion. *Bottom*: a fast-growing revertant (*Rf*), the sequencing diagram contains few overlapping peaks. *Underlined:* a base G is inserted in the deletion position and a downstream substitution (A→C). The inserted or substitutional base can be G, C, A, or T, but only one type is shown. The inserted bases of the two revertants are different because they are independent revertants. **(D)** The alignment of the wild-type, frameshifted, and repaired *bla* sequences: *bla+*: wild-type; *bla-*: frameshift; *bla*A-E*: independent revertants; *box*: the base G deleted in the OE-PCR; *underlined*: the bases inserted or deleted in the revertants; *circles*: G/C ⇌ A/T substitutions.

Competent cells of *E. coli* strain DH5*α* or BL21 were transformed with the plasmid pBR322 or pBR322-(*bla*-). The transformants were grown in tetracycline broth (TCB), 10-fold dilutions were plated on tetracycline plates (TCPs) to estimate the total number of viable bacteria and in parallel on ampicillin plates (ACPs) to test resistance. The wild type (*bla*+) grows well on both ACP and TCP (Fig 2B, left); the frameshift (*bla-*) was not expected to grow on ACPs, but a few ampicillin-resistant colonies (*bla*\*) repeatedly appeared (Fig 2B, center), which were identified as revertants whose *bla-* was repaired by reverse mutation. The possibility of carryover- or cross-contamination was excluded by blank controls (Fig 2B, right).

The screening tests were repeated 30 times by different operators, and revertants appeared in 23 out of the 30 tests. On average, the reversion rate measured in DH5*α* is 2.61×10^−8^ per cell division and reach up to 1.19×10^−7^ per cell division in BL21. Most revertants can grow in ampicillin broth (ACB), but their growth rates were much slower than wild-type *E. coli*. By subculturing at 37°C with 200 rpm shaking, it takes 36-48 hrs for most revertant to reach the late log phase while 12-24 hrs for wild-type *E. coli*. Some revertants failed to grow, but the ones survived grew faster and faster.

### 3.2 Frameshift reversion seems not to be a spontaneous mutation

Hitherto, nothing seems unusual, as the reverse mutation is a common phenomenon. It has long been assumed that reverse mutation is caused by spontaneous mutations (Fig 1): during DNA replication, spontaneous mutations occur randomly over the genome, a forward mutation causes deficient, and reverse mutations restore wild-type phenotype. If a lethal mutation occurs, most cells die, but a few survive just because they are ‘lucky’: the mutation happened to be repaired by a reverse mutation.

The classical model sounds perfect in explaining the reversion of a point mutation. However, in a ‘thought experiment’, we realized that it seems inadequate to explain the frameshift reversion observed herein.

Firstly, the reverse mutation must occur in a living cell during DNA replication or repair. Therefore, if the frameshift by itself cannot live in ampicillin, the reversion can only occur before the cells were cultured in ampicillin; otherwise, they must manage to survive in ampicillin first and repair the frameshift mutation later.

Secondly, the frameshift mutation must be repaired by inserting/deleting one (or a few more) base pairs at an appropriate location to restore the reading frame. Although small in size, a bacterial genome still contains thousands of genes, consisting of millions of base pairs. Without a mechanism for identifying and localizing the frameshifted CDS, it is extremely unlikely that a frameshift mutation can be repaired by random mutations unless a vast number of mutations are produced in the genome.

The baseline point mutation rate for wild-type *E. coli* was determined at ∼10^−3^ per genome (∼10^−9^ per nucleotide) per generation [12]. Frameshift mutation rates are about ten times lower than the baseline because InDels are rarer than base substitutions [12]. If the frameshift reversion is based on spontaneous mutagenesis, it should be even rarer and exceedingly difficult to detect. However, as mentioned above, the reversion rates of *bla*-in DH5*α* and BL21 are both hundreds of times higher than expected, suggesting that the frameshift reversion is unlikely to be caused by spontaneous mutagenesis.

Finally, point mutation is often not deleterious but tend to be neutral or beneficial; because the accumulation of point mutations brings genetic diversity, it is evolutionarily advantageous that point mutation rates are higher than reverse mutation rates. However, if frameshift mutation rates are higher than the reversion rate, the frameshift mutations accumulate in the genomes, which is unbearable. In any organism, either unicellular or multicellular, frameshift mutations are usually more dangerous than point mutations. It is unreasonable to assume that there is no mechanism to repair frameshift mutation due to the existence of multiple mechanisms for repairing point mutation.

### 3.3 The repaired gene sequences are highly divergent

To understand how *bla-* is repaired, the revertants were subcultured, and their *bla* genes were sequenced by the Sanger method. Surprisingly, in the initial revertants, their *bla* genes are often not repaired but still frameshifted (Fig 2C, top). It was puzzling how these ‘revertants’ managed to survive in ampicillin. However, as mentioned earlier, *bla-* could not be repaired if the frameshift cannot live in ampicillin. To survive in ampicillin, the frameshift may first functionalize its *bla-* gene by expressing it through translational frameshifting [13-15].

The revertants’ *bla* were sequenced for over 300 replications by Sanger sequencing, and a variety of variant *bla* sequences were obtained. In the slowly growing revertants, the sequencing diagrams of *bla* often display two sets of superimposed peaks (Fig 2C, center), indicating that they contain two *bla* genes, one frameshift (*bla-*) and the other repaired (*bla\**). The sequences of *bla\** were read from the secondary peaks by visual inspection. Some sequencing diagrams are confusing because they comprise multiple, divergent *bla\** genes. The sequencing diagrams of the fast-growing revertants have few overlapping peaks (Fig 2C, bottom) since they consist of only one or a few similar *bla\** genes. The alignment of *bla+, bla*-, and *bla\** sequences shows that insertions, deletions, and substitutions often occur at, near, or downstream, but not upstream of the base pair deleted by the OE-PCR (Fig 2D).

### 3.4 The frameshift and the revertant genome is stable

Mutator strains, such as *E. coli* mutD5, have an extraordinarily high mutation rates during DNA replication because their proofreading/mismatch repair system (PR-MRS) are defective [16]. Neither DH5α nor BL21 is a mutator. But, a strain may change into a mutator if its PR-MRS is lost or damaged during the test, a vast many mutations occur over the genome and the plasmid, which may explain the increased reversion rate.

Genome resequencing was applied to the frameshift and a revertant to test whether the strains obtained were transformed into mutators. After mapping the clean reads onto the reference genome, SNPs, InDels, and SVs were analyzed. The SNP densities are low in both the frameshift and the revertant (Table 2), indicating that neither of their PR-MRS is defective. Comparing the frameshift with the revertant, only 8 out of the 86 SNPs are different, and the remaining 78 SNP sites are shared between the two *E. coli* genomes (Supp S1), indicating that most of the SNPs are not generated during the test, but the mutations carried by the original strain.

**Table 2.** The summary of SNP/InDel variations in the two genomes

| Sample | InDels |  |  | SNPs |  |  |  |  |
| --- | --- | --- | --- | --- | --- | --- | --- | --- |
|  | INS | DEL | Total | ts | tv | ts/tv | Total | Density |
| Frameshift (*bla*-) | 2 | 4 | 6 | 52 | 30 | 1.73 | 82 | 0.01783 |
| Revertant (*bla* *) | 2 | 3 | 5 | 51 | 31 | 1.65 | 82 | 0.01783 |

Note: INS: insertion; DEL: deletion; ts: transitions; tv: transversions;

### 3.5 The variation level is higher in the repaired gene than in other genes

The other gene in plasmid pBR322, the tetracycline resistance gene (*tet*), was also Sanger sequenced to determine whether it has the same variations as *bla*. As expected, few overlapping peaks were observed in the sequencing diagrams of *tet*. By mapping the genomic and the transcriptomic reads onto the plasmid, hundreds of raw SNPs were detected in the plasmid by SNP calling, but very few passed SNP filtering (Supp S2).

DNA replication or repair, transcription, and RNA editing produce single-molecule variations (SMVs) in DNA or RNA. However, it is hard to distinguish them from noise or errors generated during NGS sequencing. In common SNP calling procedures, SNP filtering discards both sequencing errors and SMVs. So, the numbers of unfiltered raw-SNP (urSNP) were directly used to estimate the numbers of SMVs, but ambiguous loci were discarded. Due to noise and errors, it is inaccurate to estimate SMVs with urSNPs, but the differences among different genes or genotypes have true biological significance because the samples were pooled in the process of NGS sequencing, and the noise and errors should be at the same level.

The densities of urSNPs in the two genes and two genotypes are shown in Table 3. At the DNA level, the numbers of urSNPs seem exceedingly high in both genes, but urSNP densities are very low after divided by the coverage depths, indicating that the plasmid DNA replication is highly faithful. At the RNA level, although the numbers of urSNPs are much smaller than those at the DNA level, the urSNP densities are much higher than those at the DNA level, considering the depths of coverage.

**Table 3.** The count and density of raw SNPs of the two genes in the plasmid pBR322

| Sample | Average<br>depth<br>of coverage | Total<br>(4,362bp) |  | *Tet*<br>86-1276<br>(1191 bp) |  | *bla*<br>3293-4153<br>(861 bp) |  |
| --- | --- | --- | --- | --- | --- | --- | --- |
|  |  | Count | Density | Count | Density | Count | Density |
| *Frameshift-DNA* | 7,859 | 292 | 0.0083 | 76 | 0.0081 | 61 | 0.0090 |
| *Revertant-DNA* | 7,826 | 298 | 0.0085 | 70 | 0.0075 | 67 | 0.0099 |
| *Wildtype-RNA* | 188 | 18 | 0.0213 | 5 | 0.0224 | 2 | 0.0124 |
| *Revertant-RNA* | 181 | 25 | 0.0307 | 6 | 0.0278 | 6 | 0.0385 |

At the DNA level, the urSNP density of *bla* is slightly higher than that of *tet* in the frameshift but much higher in the revertant. At the RNA level, the differences between the two genes/genotypes are even more obvious than those at the DNA level. The urSNP density of *bla* is lower than that of *tet* in *Wt* but higher than that of *tet* in *Rs*. In summary, significantly more urSNPs are observed in repaired genes than in wild-type genes (*bla* or *tet*) at both DNA and RNA levels.

Besides, if the *bla\** variations were caused by random mutagenesis, the number of variations in *bla* should be proportional to those in the plasmids and the genomes. Thus, the number of SNPs of the genome or plasmid in the revertant should be much higher than those in the frameshift. However, the numbers of SNPs are comparable in the two genomes (Table 2), or plasmids (Table 3). In other words, the number of *bla* variations is independent of those of the plasmid or the genome, suggesting that the high-level of *bla* variations are not caused by random mutagenesis.

### 3.6 PTC signals the repair of the frameshift mutation

Above analysis shows that the reversion of *bla*-seems not to be random mutations, then it must be a targeted gene repair. The nonsense mRNAs transcribed from *bla*-must be involved in the repair since the frameshift mutation is not recognizable in the double-stranded DNA. In principle, PTC is the only signal that can be recognized in nonsense mRNAs. Consequently, if no PTC is present, a nonsense mRNA cannot be recognized, even if its reading frame is wrong.

To prove that PTC signals frame repair, a *PTC-substituted bla-* gene (*bla*#) was designed by replacing each PTC in *bla*-with a sense codon in line with the readthrough rules derived from *E. coli* suppressor tRNAs (Table 1). The *bla*# gene was synthesized, cloned into the vector pET-28a, and expressed in *E. coli* strain BL21. The transformants were plated on ACPs to screen for revertants. Their *bla* genes were Sanger sequenced. As expected, no revertant was detected by screening tests, and there are few overlapping peaks in their sequencing diagrams, indicating that *bla#* was not repaired.

SDS-PAGE of the protein extract shows a protein band of the predicted molecular weight of BLA# (33.85kDa), representing the *bla*# product (Fig 4A). The BLA# protein was purified by HisTrap™ HP and showed a single band for the purified BLA# in the SDS-PAGE (Fig 4B). No lactamase activity was detected by the iodimetry test of the product (Fig 4C). The expression of a defective BLA# confirms that *bla*# is not repaired as *bla*-but is expressed in the wrong reading frame.

**Fig 3.**
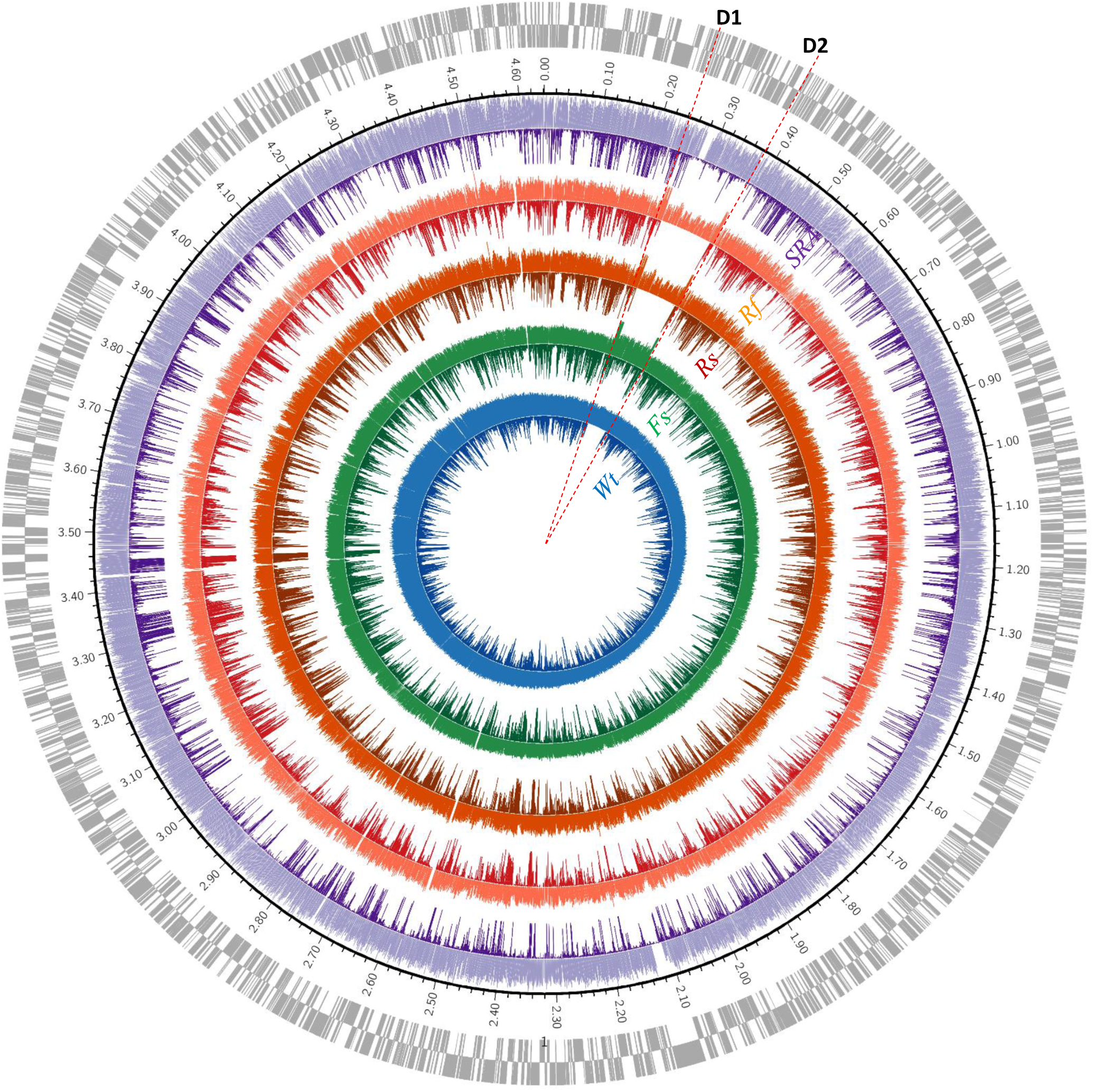
The circular map shows the depths of coverage of the genomes and the transcriptomes. Circos was used to display the depth of coverage of each sample or dataset. The circle outside of the coordinate circle is the annotated CDSs in the sense and antisense strand; the circles inside the coordinate circle are the depth of coverages for transcriptomic and genomic reads. *Blue* (*Wt*): the wild type; *green* (*Fs*): the frameshift; *red* (*Rs*): the slow-growing revertant; *orange* (*Rf*): the fast-growing revertant; *purple* (*SRA*): the SRA dataset; *inward:* transcriptomes; *outward*: genomes.

**Fig 4.**
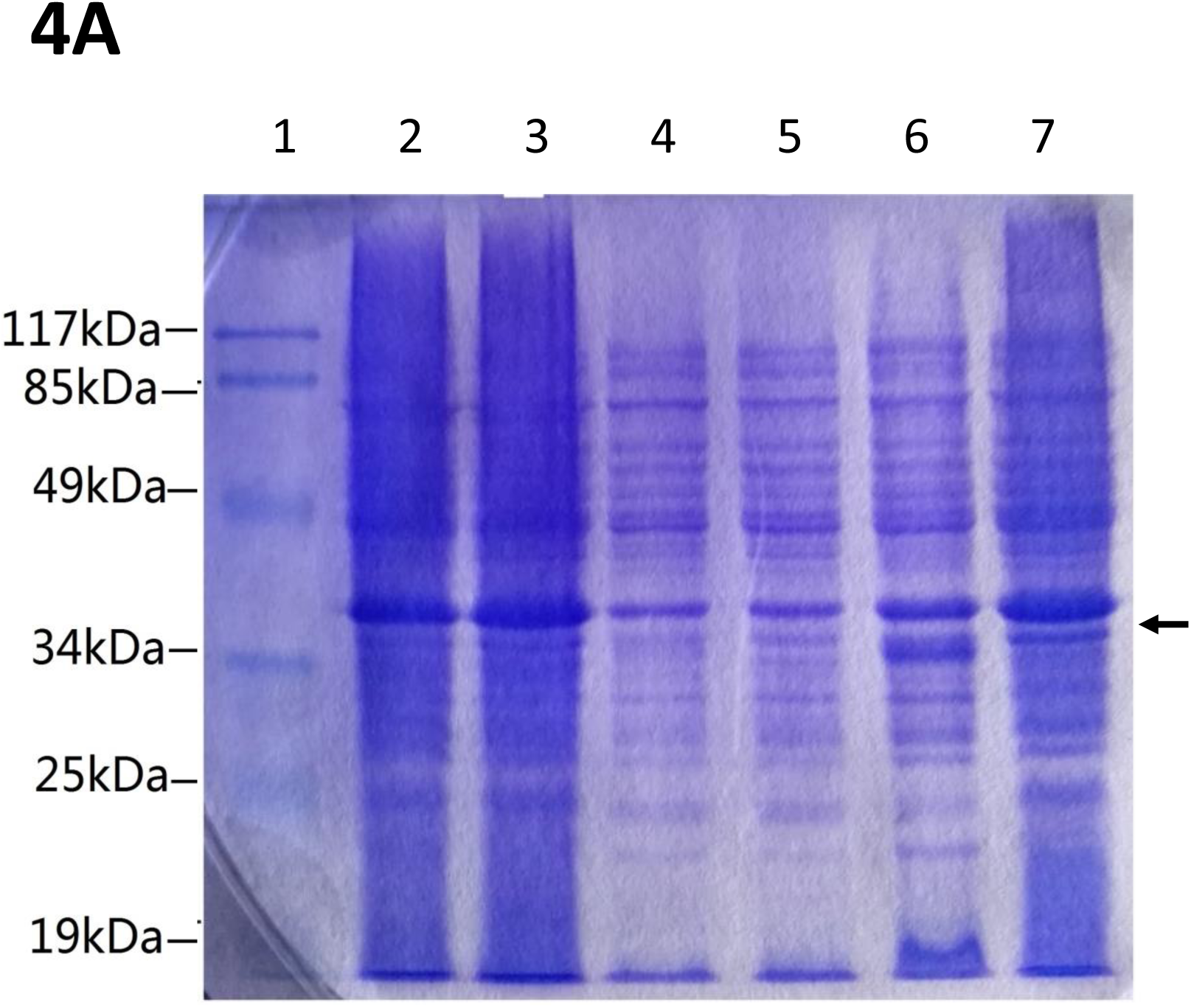
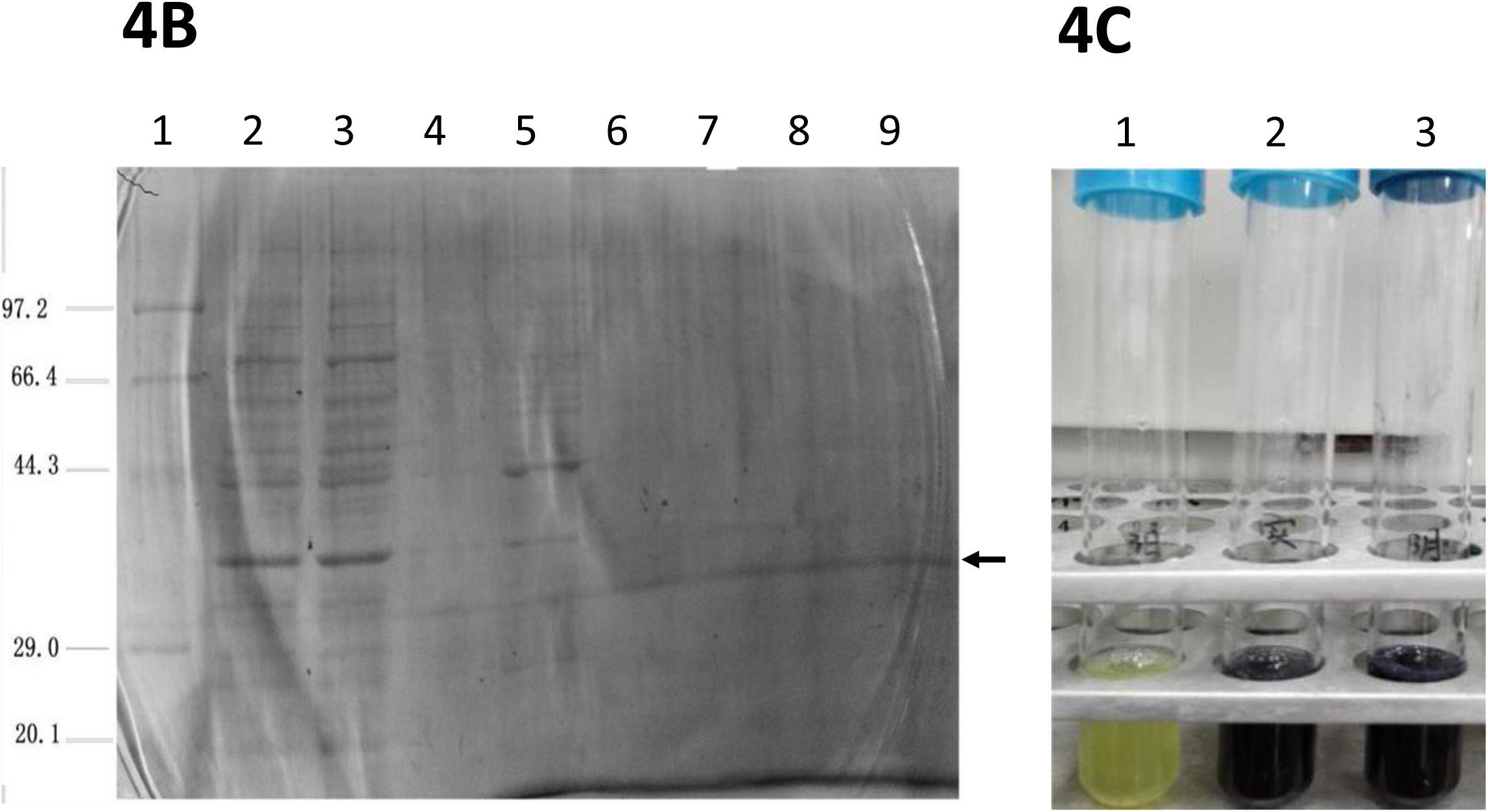
Expression of the PTC-substituted frameshift *bla* (*bla*#) in *E. coli* BL21. **(A)** SDS-PAGE of the lysates of pET28a-(*bla#*) recombinant *E. coli* cells; lane 1: the protein marker; lane 2 and 3: the whole-cell lysates of uninduced and induced cells; lane 4 and 5: the supernatant of uninduced and induced cells; lane 6 and 7: the precipitate of uninduced and induced cells; **(B)** Purification of the BLA# protein by the nickel column chromatography. Lane 1: the protein marker; lane 2: the supernatant of induced cells; lane 3: the outflow of the nickel column; lane 4: the discharge of the washing buffer; lane 5-9: different elution components. **(C)** Detection of β-lactamase activity by iodimetry: **(1)** the revertants of *E. coli* BL21 with pET28a-(bla-) causes the fading of the iodine/starch solution, suggests that *bla-* is repaired, producing an active β-lactamase, transforms ampicillin into penicillium thiazole acid, which binds with starch and competitively inhibits the binding of iodine to starch; **(2)** *E. coli* BL21 with pET28a-(bla#) does not cause the fading, suggesting that the *bla#* gene is not repaired but expressed an inactive product; **(3)** the negative control, the empty *E. coli* BL21, with no plasmid, no *bla*, no β-lactamase, no fading.

The major difference between the two *bla* genes is that *bla-* contains PTCs while *bla*# does not. Therefore, the PTCs must signal the frame repair since they are the only possible signal for the cells to recognize the nonsense mRNA through surveillance. The two *bla* may be different in other respects, such as GC contents and secondary structures of encoded RNAs, but these characteristics are unlikely to play any role in frame repair because they are not useful for recognizing nonsense mRNAs.

### 3.7 Regulation of DNA and RNA manipulating genes in frame repair

The transcription levels for all genes are profiled by RNA sequencing for four *E. coli* strains, including *Wt, Fs, Rs*, and *Rf*. The *E. coli* genome was annotated with 4814 genes, including protein-coding genes, mobile elements, tRNAs, rRNAs, non-coding RNAs, and pseudogenes. In the transcriptome analysis, 4347 known and 16 novel genes were detected (Supp S3). Hundreds of genes show great changes (*q* < 0.005 and *f* ≥ 2.0, up or down) in their transcription levels in *Fs, Rs*, and *Rf* compared to *Wt* (Supp S4). Enrichment analysis identified dozens of KEGG pathways associated with the DEGs (Supp S5). However, most of these pathways are not related to gene repair but nutrition metabolism and ampicillin resistance due to the different media components.

Frame repair is a complex process. Many pathways, such as transporting, localizing, and metabolizing nucleotides, amino acids, and other nutrition, may play roles in this process. However, the key genes should directly manipulate genetic materials (DNA or RNA). Therefore, here we focus on the KEGG pathways that directly manipulate DNA or RNA, including DNA replication, transcription, base or nucleotide excision repair, mismatch repair, homologous recombination, RNA degradation, RNA processing, RNA editing, translation, and tRNA synthesis.

By comparing *Fs, Rs*, or *Rf* with *Wt*, the transcription of DNA/RNA manipulating genes show 28 great changes, 26 up and 2 down (Table S1). When moderate changes (*f* ≥1.2 and *q* < 0.05) were considered, they show 284 changes, 219 up and 65 down (Table S1). These changes revealed how genes are regulated in the process of frame repair:

First, in most DNA or RNA manipulating pathways (Table S2-S11), except for the base excision repair, the numbers of up changes are greater than those of down changes; in the translation pathway (Table S12), however, the number of down changes is much greater than that of up changes (Fig 5A). Specifically, genes encoding the core subunits of DNA polymerase III (Fig 5B), RNA polymerase (Fig 5C), RNA degradosomes (Fig 5D), tRNA synthetases (Table S10), and RNA processing proteins (Table S11) were all or mostly upregulated in *Fs, Rs* and *Rf*. However, most ribosomal proteins were reduced in *Fs* and *Rs* but rose in *Rf* (Fig 5E, Table S12). In *Rs*, DNA replication, transcription, RNA degradation, and RNA processing are active, while protein biosynthesis is reduced, indicating that cells are engaged in gene repairing, leading to their slow growth. In *Rf*, however, the protein biosynthesis and the cell growth recovered because its *bla* gene is less problematic, and thus, gene repair requires less cell energy.

**Fig 5.**
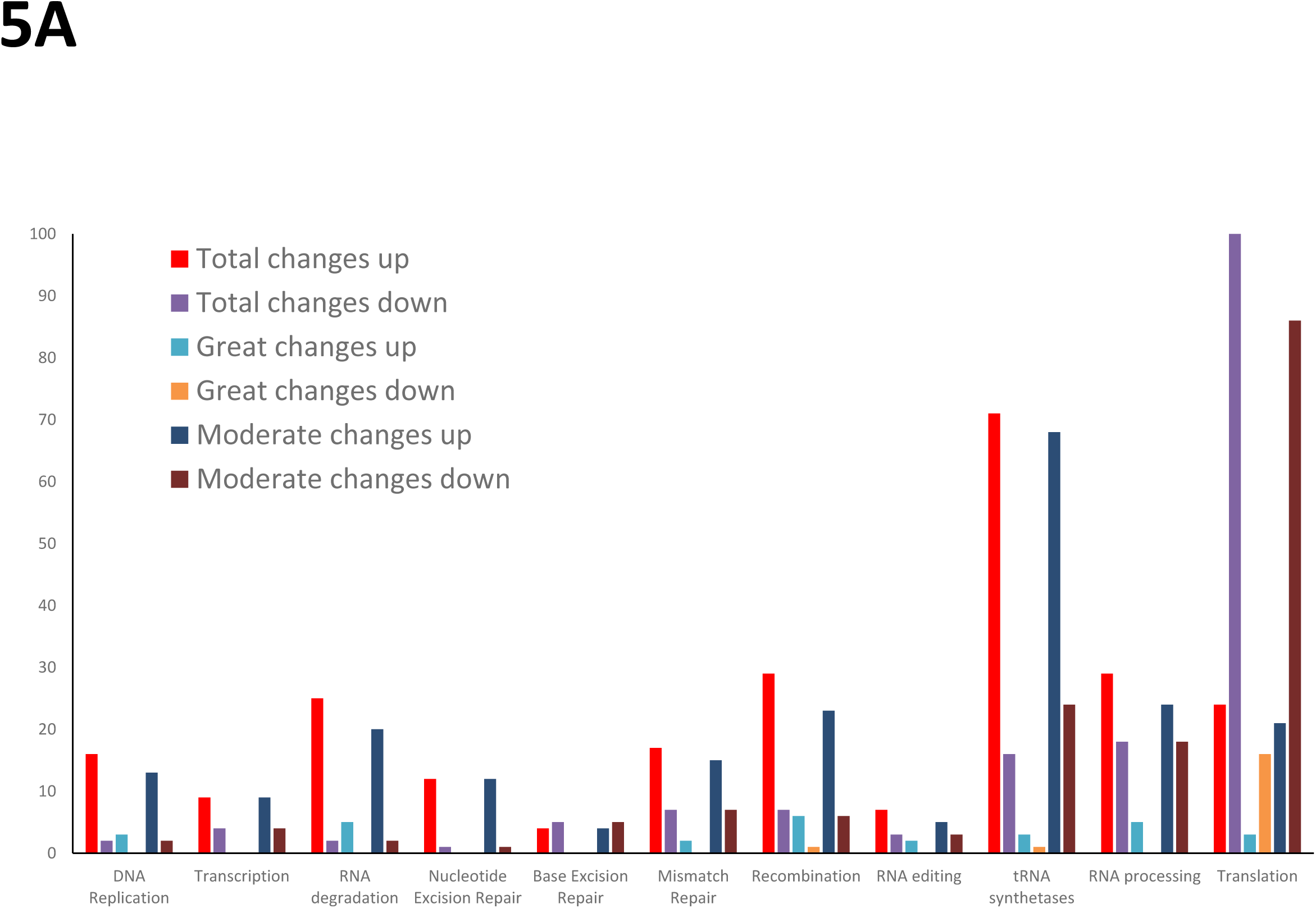
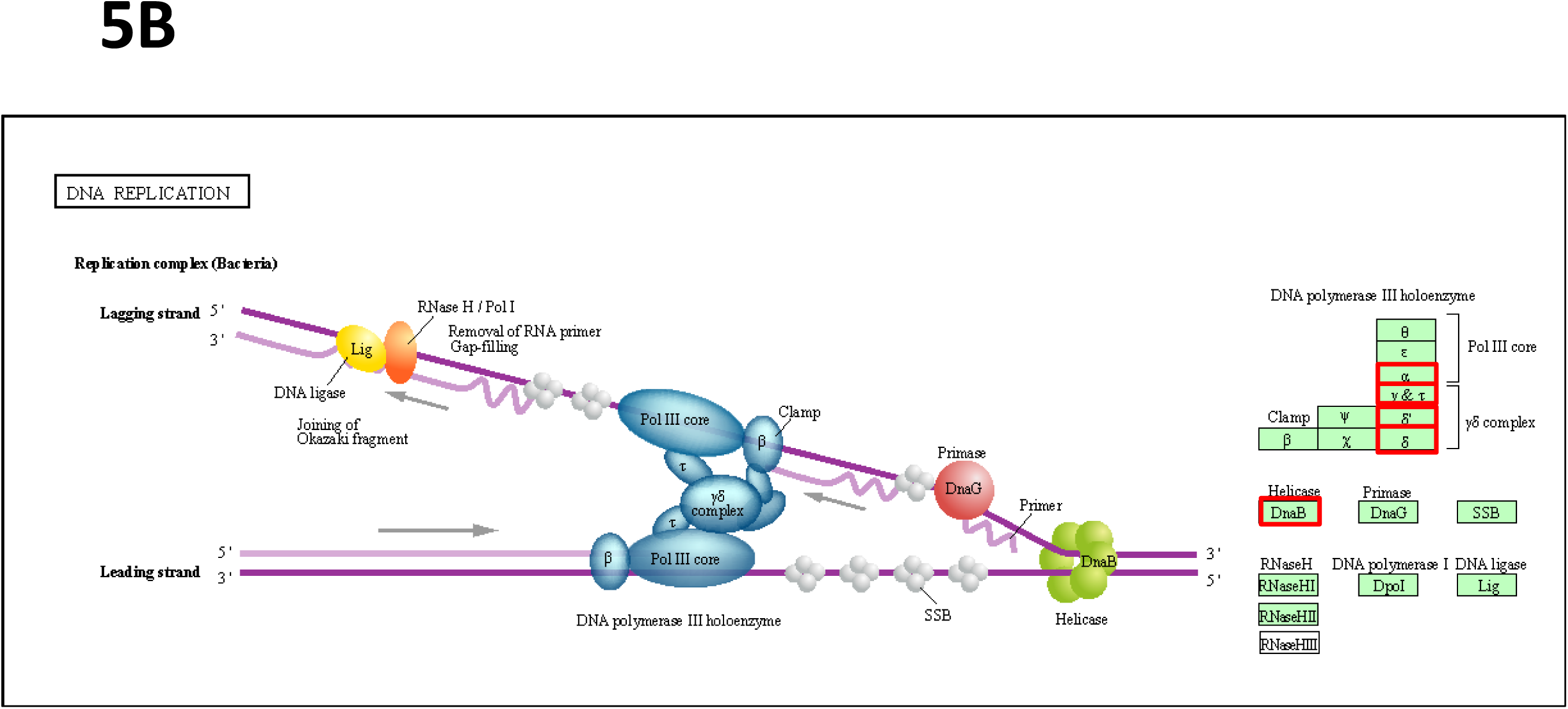
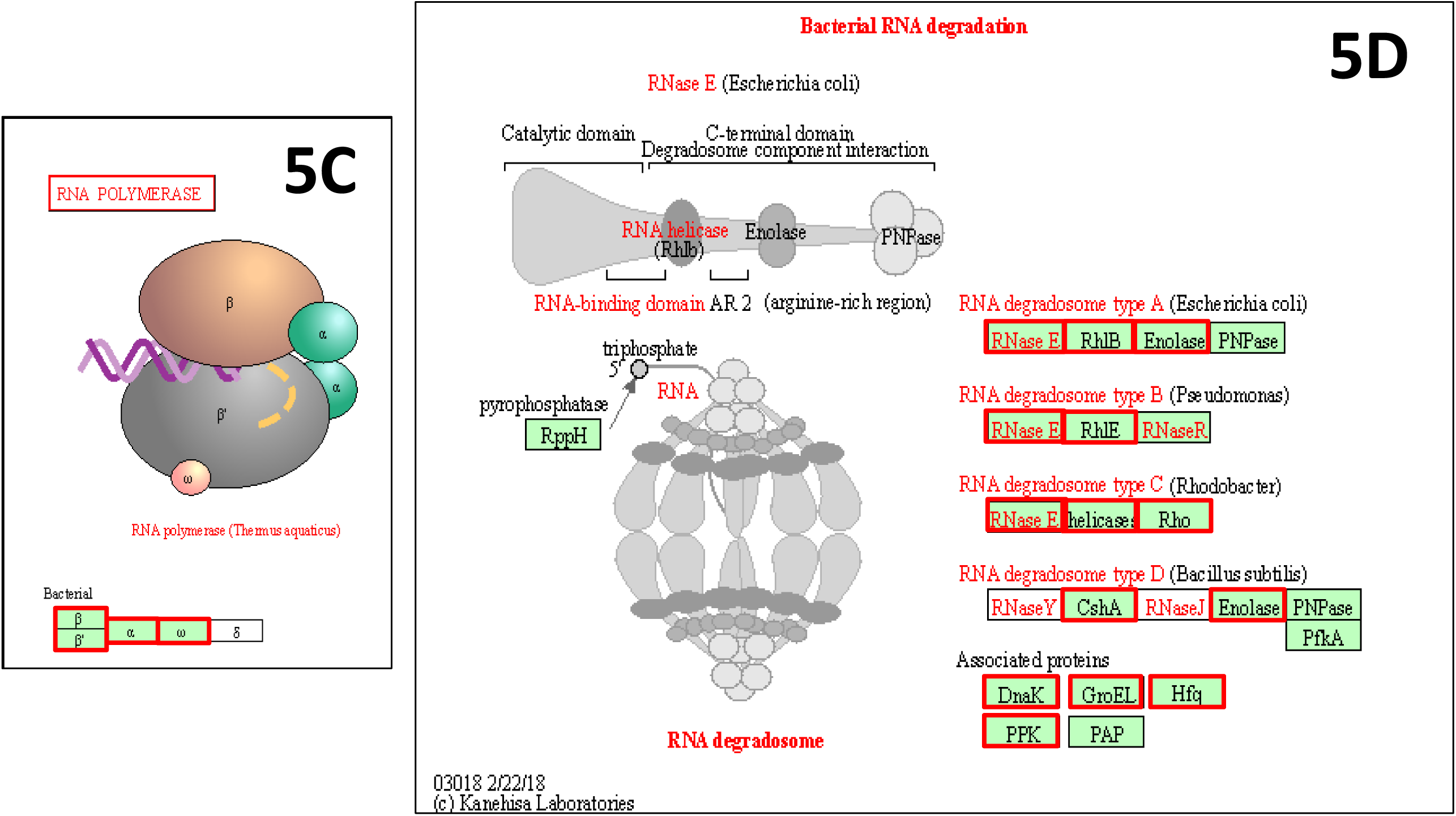
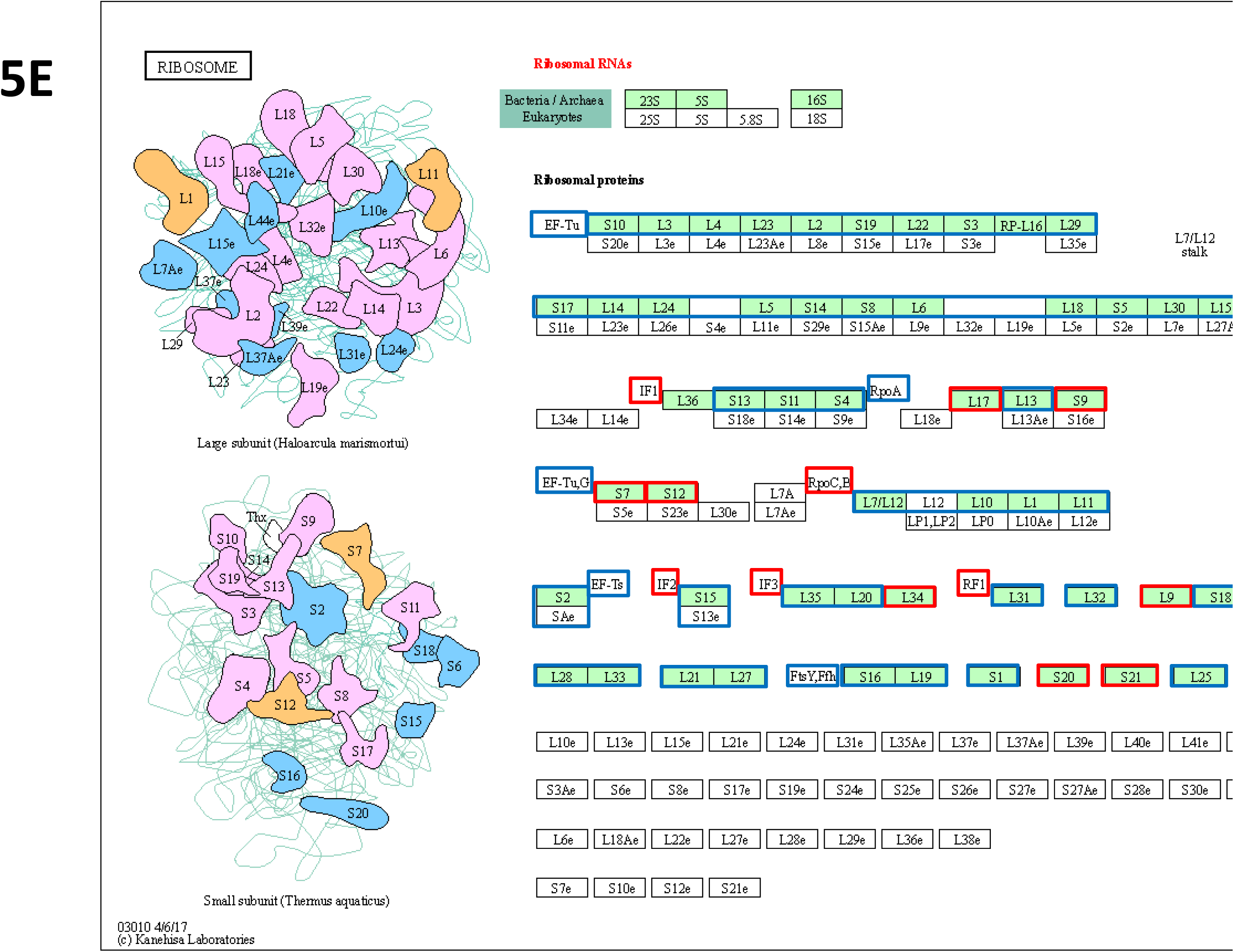
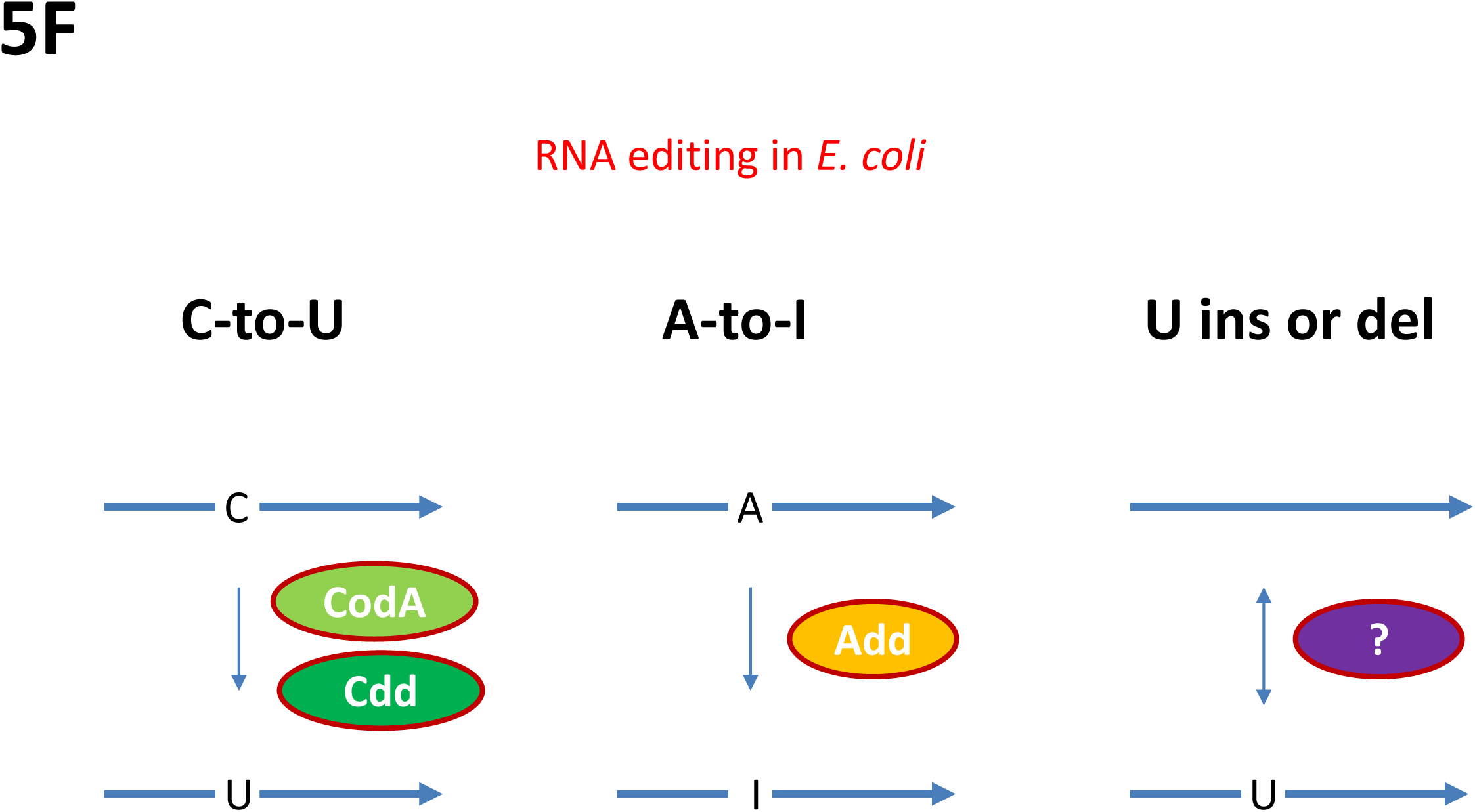
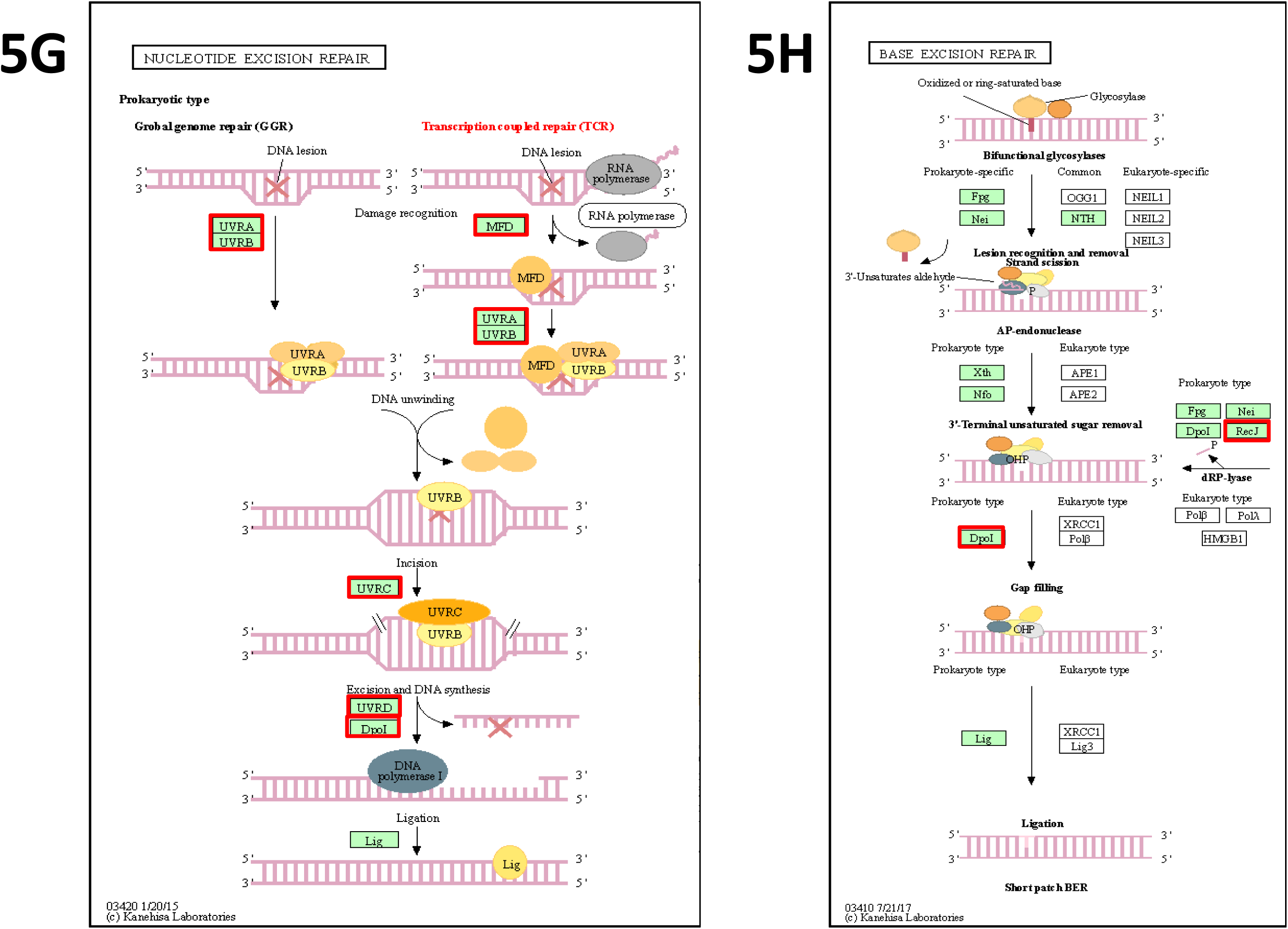
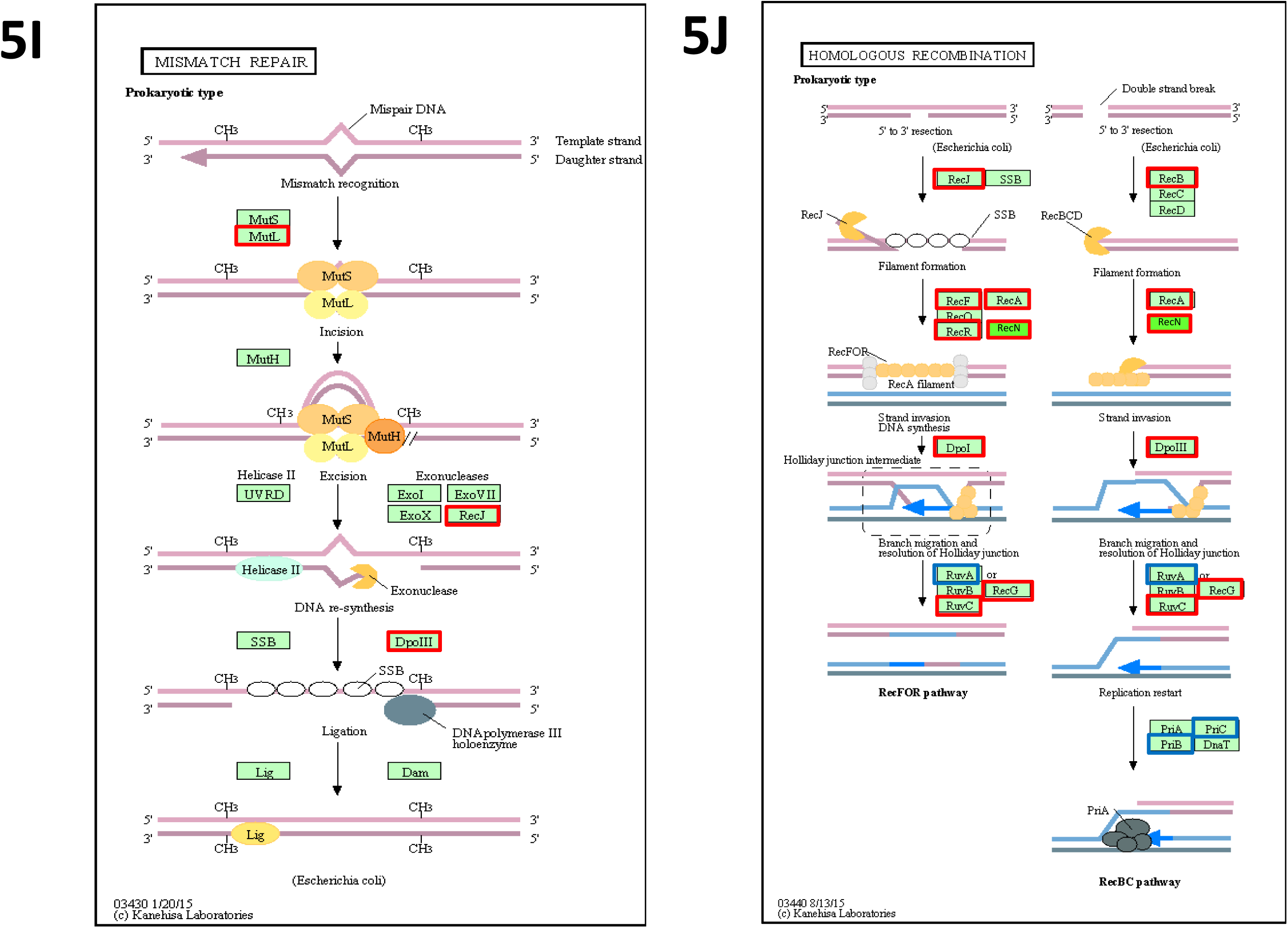
DEGs identified in the DNA and RNA manipulating pathways by the transcriptome analysis. **(A)** the number of changes in the DNA or RNA manipulating genes by comparing *Fs, Rs, Rf* with *Wt* (Table S1), and the changes in **(B)** DNA replication (Table S2), **(C)** RNA polymerase (Table S3), **(D)** RNA degradation (Table S4), **(E)** translation (Table S12), **(F)** RNA editing (Table S9), **(G)** nucleotide excision repair (Table S5), **(H)** base excision repair (Table S6), **(I)** mismatch repair (Table S7), **(J)** homologous recombination (Table S8); *double/single lines*: DNA/RNA; *color shapes:* genes or proteins; *red/blue border*: upregulated/downregulated in *Fs, Rs*, or *Rf* compared to *Wt*.

Second, the cytosine or isoguanine deaminase gene (*codA*) and a putative metallo-dependent hydrolase domain deaminase gene (*yahJ*) are expressed only in *Fs* but not in *Wt, Rs*, or *Rf*; the cytidine/deoxycytidine deaminase gene (*cdd*) is upregulated in *Rf*, and the adenosine deaminase gene (*add*) rose in both *Fs* and *Rs*. These deaminases are all responsible for RNA editing. CodA and Cdd catalyze C-to-U editing, while Add for A-to-I (Fig 5F). Inosine (I) is structurally similar to guanine (G), leading to I to cytosine (C) binding [17]. In contrast, the tRNA-specific adenosine deaminase gene (*tadA*) is not changed among *Wt, Fs, Rs*, and *Rf* (Table S9), which is consistent with the fact that TadA is responsible for the A-to-I editing at the wobble position 34 of tRNA-Arg2 [18].

Since C-to-U and A-to-I editing produce G/C ⇌ A/T substitutions, but not G ⇌ C or A ⇌ T, we classified base substitutions into two subtypes: G/C ⇌ A/T as type A; G ⇌ C and A ⇌ T as type B. Sanger sequencing show that type A substitutions occur more frequently than type B in *bla\** (Fig 2D, circles). The same bias was also observed in the urSNPs of the revertant’s plasmid (Table 4). The nA/nB ratio indicates that type A SNPs appear more frequently than type B in both genes, and it is more biased to type A in *bla\** than in *tet*, indicating that the urSNPs of *bla*\* are likely derived from RNA editing.

**Table 4.**
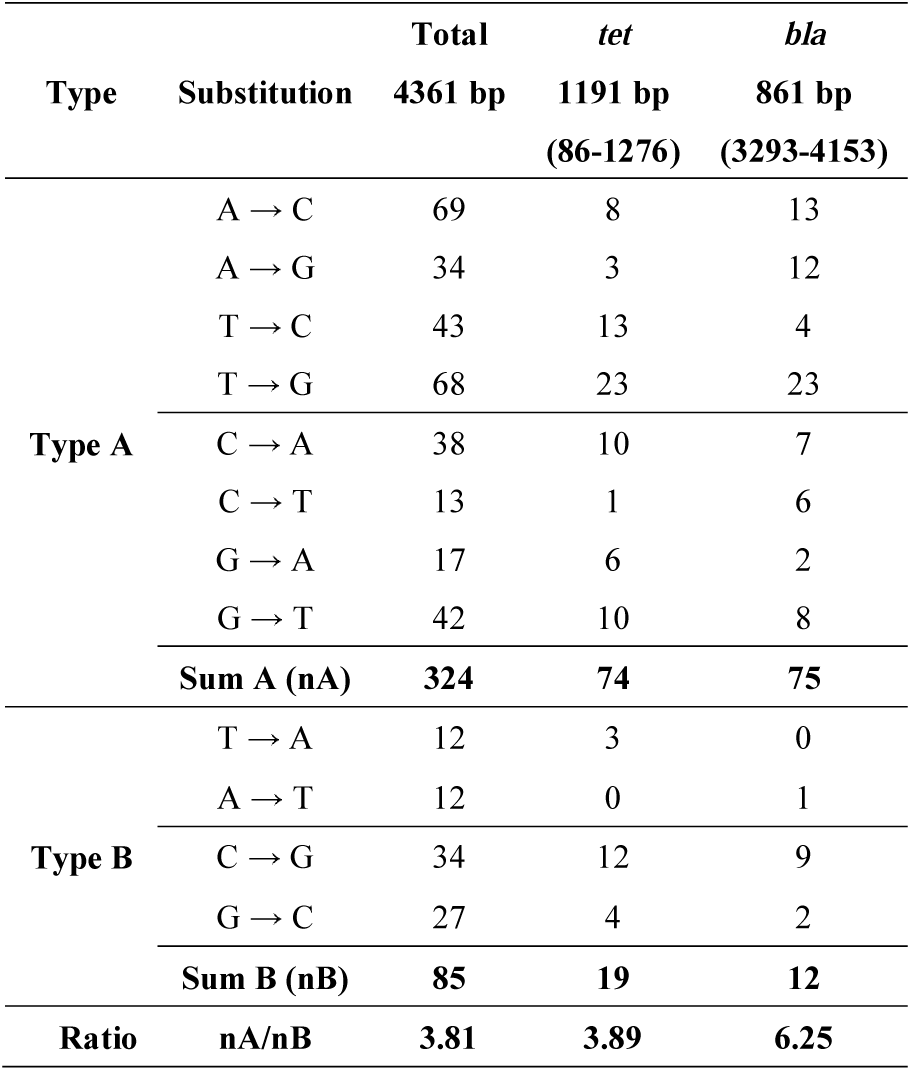
The two different types of raw SNPs in the two genes of the revertant’s plasmid

Third, in *Fs, Rs*, and *Rf*, the nucleotide excision repair pathway is upregulated (Fig 5G, Table S5), but not the base excision repair pathway (Fig 5H, Table S6), reasonably indicating that frame repair is achieved by nucleotide excision rather than base excision. However, in the methyl-directed mismatch repair pathway, neither MutS nor MutH is differential, and MutL is upregulated (Fig 5I), but the G/U mismatch-specific/xanthine DNA glycosylase (Mug) is downregulated in all *Fs, Rs*, and *Rf* (Table S7). RNA editing produces substitutions and insertions of uracil base in the edited mRNAs. Consequently, there must be many G/U mismatches between the template mRNA and the target CDS. Usually, G/U mismatches are proofread into G:C base pairs by mismatch repair or base excision repair [19]. In frame repair, however, G/U mismatches might be converted into A:T base pairs instead because the CDS is repaired using an mRNA template, and both the base excision repair and the G/U mismatch-specific proofreading are downregulated.

Fourth, the recombination and repair (Rec) proteins, including RecA, RecB, RecJ, RecF, RecG, RecN, RecR, and RuvC, are upregulated in *Fs* and *Rf* (Fig 5J), suggesting that they might be involved in the process. As mentioned earlier, the reversion rate of BL21 (with a functional RecA) is 4-5 times higher than that of DH5*α* (with a defective RecA). Recently, it was found that RecN functions in the strand invasion step of RecA-mediated DNA recombination [20]. The Rec proteins are upregulated in *Rf* but not in *Rs*, suggesting that they may play a dual role in frame repair: on the one hand, DNA recombination may interfere with RNA-directed DNA repair, which is adverse to frame repair; on the other hand, recombination among different *bla*\* can promote cell survival and growth. In *Fs* and *Rs*, the main task is to repair the wrong reading frame of *bla*, not by DNA recombination but by RNA-directed DNA repair, nucleotide excision repair, and mismatch repair. In *Rf*, however, the *bla* reading frame may have been repaired, but the repaired genes consist of a variety of variations. By upregulating the Rec proteins, recombination among different *bla*\* genes improves cell survival and growth.

Finally, comparing with *Wt*, the elongation factors, EF-Tu 1, EF-Tu 2, and EF-Ts, reduced in *Rs* but rose in *Rf*, and the selenocysteinyl-specific elongation factor (SelB) increased in *Rf* (Table S12). SelB is a specialized elongation factor that replaces EF-Tu to insert selenocysteine into a peptide chain in the place of codon UGA [21, 22]. The regulation of these elongation factors indicates that in *Rf*, the protein biosynthesis is restored, and the remaining stop codons in mRNAs can be readthrough.

### 3.8 Silent genes and transposons activated in the frameshift

By plotting the coverage depths of all genomic and transcriptomic datasets on the same circular map (Fig 3), including *Wt, Fs, Rs, Rf*, and the pooled SRA dataset (*SRA*), it is shown that the genomes of *Fs, Rs* and *Rf* harbor a duplicate region (*D*) from 0.24 to 0.38 MB, showing higher coverage depths than the other genomic regions, especially at the ends (D1 and D2). Surprisingly, genes in the center of this region are completely silenced in *Wt, Rs* and *Rf* but actively transcribed in *Fs*. In *SRA*, the transcription levels in this region are consistent with those of *Wt*, except for a few genes peak in the center, confirmed that most genes are either silenced or transcribed at low levels in the center of this region in wild-type *E. coli*.

Extensive protein occupancy domains (EPODs) have been discovered in *E. coli* by using an *in vivo* protein occupancy display (IPOD) technology [23]. Interestingly, the majority of EPODs are localized to transcriptionally silent loci dominated by conserved ORFs. YafF (coordinate 236644-239264), a putative uncharacterized protein identified in a transcriptionally silent EPOD (tsEPOD), locates just at the start of this region (D1), implicating that this region is the organizing center for a topologically isolate domain [23], harboring specific, functionally important genes.

In the annotation file for *E. coli*, this region is annotated with 149 genes (Supp S6). Interestingly, the left end (D1) mainly includes genes for DNA replication, DNA repair, transcription, translation, and RNA processing, including the DNA polymerase IV (*dinB*), the ribosome-dependent mRNA interferase (*yafO* and *yafQ*), the peptide chain release factor (*prf*), and the RNA polymerase holoenzyme assembly factor (*crl*). Especially, DNA polymerase IV is error-prone because it exhibits no 3′→5′ exonuclease (proofreading) activity and is responsible for non-targeted mutagenesis and SOS DNA repair. Pol IV is switched on in SOS DNA repair caused by stalled polymerases at the replication fork in *E. coli* [24]. Pol IV transcripts were greatly upregulated in *Fs, Rs* and *Rf* (Table S2), indicating that frame repair might require Pol IV and thus may connect with non-targeted mutagenesis and SOS DNA repair; but it does not mean that frame repair is just non-targeted mutagenesis or SOS DNA repair because, as above described, it does not lead to a great many of random mutations throughout the whole genome but targeted specifically to the frameshift gene.

Interestingly, the center of this region contains three genes that encode the cytosine or isoguanine deaminase (CodA), cytosine transporter (CodB), and a putative metallo-dependent hydrolase domain deaminase (YahJ). As mentioned earlier, these genes are expressed only in *Fs* but not in *Wt, Rs* or *Rf*. As *yahJ* and *codA* are regulated consistently (Table S9), it seems that they belong to the same operon. Because CodA is responsible for C-to-U editing, YahJ may also play a role in RNA editing or frame repair.

Besides, the center of this region harbors many transposons, including the inserted sequences (IS1, IS2, IS3, IS5, IS30) and the prophage genes (transposase, integrase, and resolvase). As is well known, transposable elements are rich in repetitive sequences, which may cause higher coverages in this region than in the other regions. However, it was puzzling that the ‘duplicate region’ causes higher coverages only in *Fs, Rs* and *Rf* but not in *Wt* (Fig 3). Presumably, in *Wt*, the central genes and the transposons are all silenced; in *Fs*, the transcribing of the central genes leads to the activation, proliferation, and translocation of the transposons. Consequently, NGS produces higher coverages in this region, especially at the two ends, than in the other genomic regions. In *Rs* and *Rf*, however, the central genes and the transposons were re-suppressed, and high coverages are produced only at the ends. The structural variations found in this region (Supp S7), such as Δ257911-258669 and Δ257906-258670, validated this assumption because they contain some of these transposons and appear in the genomes of both *Fs* and *Rs*.

### 3.9 Detection of the gene products by proteome profiles

The total protein samples of *Fs* and *Rs* were analyzed by a quantitative analysis of the global proteome. In total, 2,029 proteins were identified, among which 1,706 proteins were quantified. The mass error of the peptides is around 0.0. The length of the peptides is distributed mostly between 8 and 16. GO annotations of the identified peptides were derived from the UniProt-GOA database or InterProScan.

Post-transcriptional processing plays a critical role in regulating protein levels and leads to discordance between protein and transcript levels [25]. Therefore, the proteome profiles are used not to identify differentially expressed proteins but to verify the above gene products. As highlighted in red in Supp S8, the identified peptides confirmed the expression of the DEGs identified, including the cytidine deaminase (Cdd), the cytosine deaminase (CodA), the adenosine deaminase (Add), the mRNA interferase (RelE), the transposases (InsC, InsE, InsL), the G/U mismatch-specific DNA glycosylase (Mug), and the selenocysteine-specific elongation factor (SelB), and so on.

The proteome analysis also confirmed the presence of many other proteins that are involved in RNA editing, RNA processing, or DNA repair, such as the endonuclease V (EndoV), the DNA recombination protein (RmuC), the DNA mismatch repair protein (MutS/MutL), and the DNA-damage-inducible protein I (DinI), and so on. Their protein and transcript profiles show no difference among the 4 different genotypes, but it does not mean that they must not involve in frame repair, because housekeeping genes may also play roles in the frame repair process.

## 4. Discussion

### 4.1 A new model for frame repair

In the above, we demonstrated that PTC signals nonsense mRNA recognition and subsequent frame repair. Sanger and NGS sequencing suggest that the substitutions in *bla\** are derived from RNA editing. The transcriptomes show that many DNA and RNA manipulating pathways are involved. Based on these data, we propose a six-step model for frame repair (Fig 6), referred to as *nonsense-mediated gene revising* (*NMGR*):

**Fig 6.**
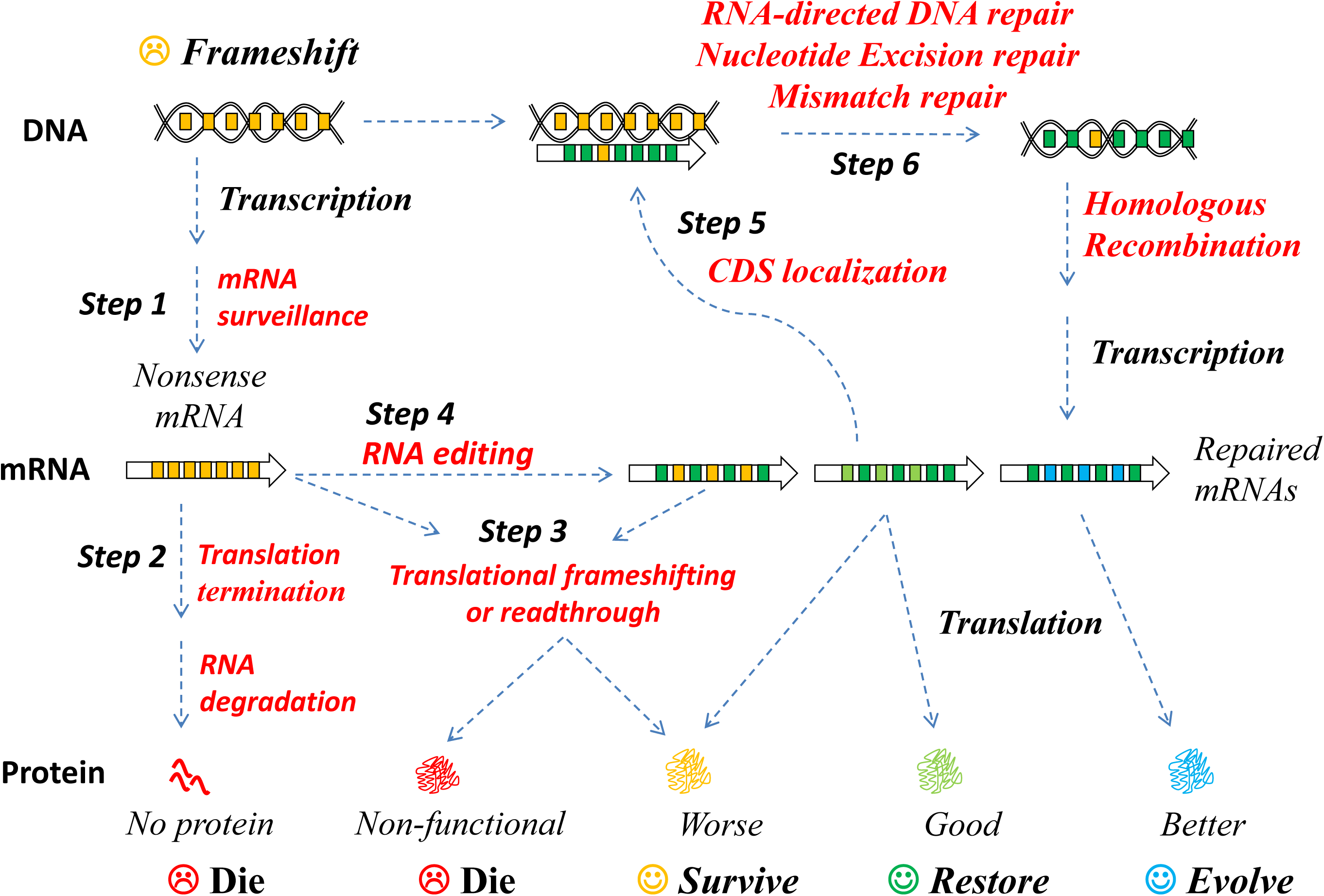
A new model for frame repair: nonsense-mediated gene revising (NMGR). *Step 1*: a frameshift gene is transcribed and translated. The nonsense mRNAs containing many PTCs (*yellow bars*) are identified through mRNA surveillance by recognizing the PTCs; *Step 2*: nonsense mRNAs cause *translational termination* and are degraded by NMD; *Step 3*: nonsense mRNAs are translated by translational frameshifting or readthrough; *Step 4*: nonsense mRNAs are repaired by editing, PTCs are substituted (*green bars*) through base pair insertion, deletion, and substitution; *Step 5*: a repaired mRNA is transported to identify and localize the defective CDS; *Step 6*: the frameshift gene is repaired using the edited mRNA template, then transcription and translation resume, producing mRNAs and proteins. The repaired gene sequences are usually different from the original sequence, the remaining stop codons are readthrough, and the process repeats until the gene is restored, and the host recovered and sometimes evolved.

**Step 1. *Nonsense mRNA recognition***: When a frameshift gene is transcribed, the nonsense mRNAs are recognized by mRNA surveillance [26]. A ribosome is blocked when it encounters a PTC while translating a nonsense mRNA [27]. Then, the nonsense mRNAs are subjected to either one of the following pathways, degradation (step 2), translation (step 3), or editing (step 4).

**Step 2. *Nonsense mRNA degradation***: nonsense mRNAs are degraded through nonsense-mediated mRNA decay (NMD) in eukaryotes. It is unclear how nonsense mRNAs are processed in bacteria, but their mRNA quality is also tightly controlled [28]. NMD may also exist in bacteria though it has not been characterized.

**Step 3. *Nonsense mRNA translation*:** Nonsense mRNAs are not always degraded but sometimes translated through translational frameshifting or readthrough. If the gene is essential, then a cell survives if sufficient proteins are produced or die if not.

**Step 4. *RNA editing***: The nonsense mRNAs must be used for the recognition and repair of the frameshift gene, but their sequences are the same as the frameshifted CDS, which are defective by themselves and not directly useful for templating DNA repair. Therefore, the nonsense mRNAs are edited before they are used to template the repair.

***Step 5. CDS localization:*** After editing, if the translation resumes the production of proteins, the repaired mRNAs are transported back to the genomic or plasmid DNA to localize its defective CDS.

**Step 6. *RNA-directed DNA repair***: The CDS is repaired using a functional mRNA as a template by RNA-directed DNA repair, mismatch repair, and nucleotide excision repair. Homologous recombination may also involve in a later stage of this process. The cell recovers if sufficient functional mRNAs and proteins are produced by the standard transcription and translation.

### 4.2 NMGR explains frameshift reversion and the variations of the revertant

As mentioned above, the *bla\** sequences are usually different from the original *bla*, and most InDels and substitutions are located at, near, or downstream of the frameshift-causing deletion, but not upstream (Fig 2C-2D), indicating that the PTCs not only signal the recognition of the nonsense mRNAs but also serve as flags for RNA editing. Since the PTCs appear only downstream of the deletion, RNA editing and subsequent DNA sequence changes are also restricted to the same region.

When cells are cultured in an ampicillin medium, the survival of a cell depends on if it can produce sufficient many functional proteins. Each CDS can transcribe hunreds to thousands of many mRNAs, each can be edited differently without changing the CDS. Consequently, the probability of producing a functional mRNA through mRNA editing is much greater than that of direct random mutation in genomic DNA.

### 4.3 Previous studies are supportive of NMGR

The NMGR model integrates several key links, including RNA surveillance, RNA editing, DNA repair, and recombination, all of which have been intensively studied. If this mechanism exists, like the connected pathways, it should be widely existing and highly conserved among species. In the supplementary text, we present a review of previous studies on these links: (1) mRNA decay is linked to other nonsense mRNA processing pathways; (2) RNA can direct DNA repair; and (3) RNA editing is linked to DNA repair. All in all, the crosslinking of these pathways forms a network supporting the NMGR model.

In conclusion, it is shown that PTC signals frame repair and the nonsense mRNAs are edited before directing CDS repair. Thanks to NMGR, the DNA-level mutation rate keeps low but temporarily rises in a particular gene at the RNA level. An RNA-level mutation is preserved in DNA if it is advantageous. If an mRNA or a protein is better optimized in a cell, it is preserved in the genome so that the cell and its progeny obtain better adaptability and evolutionary advantage over their competitors. Thereof, NMGR provides an ingenious solution to the paradox between the mutation rate and the power of selection, accelerating evolution progress without the cost of a high mutation rate in the genome. While repairing a frameshift muation, NMGR brings diversity to the gene and surviving potential to the cells, acting as a driving force for molecular evolution.

Finally, we show that frame repair is a complex problem involving hundreds of genes, many of which are uncharacterized, and many problems are far from clear. In future studies, it is necessary to elucidate the process in *E. coli* and other organisms, especially multicellular, eukaryotic species.

## Data accessibility

The preprint version of this article and a supplementary text file are available from bioRxiv at https://doi.org/10.1101/069971, including a detailed materials and methods, a review of the previous stydies, references, and Table S1-S12. Supplementary data files are available from FigShare at https://doi.org/10.6084/m9.figshare.11497128.v5, including **S1**: the list of SNPs in the genomes; **S2**: the list of SNPs in the plasmid; **S3**: the read count and FPKM for all genes; **S4**: the list of DEGs; **S5**: the list of enriched KEGG pathways; **S6**: the list of genes annotated in the duplicate region; **S7**: the list of structural variations; **S8**: the list of peptides identified in the proteomes.

## Author Contributions

Xiaolong Wang conceived the study, designed the experiments, analyzed the data, plotted the figures, and wrote this paper. Xuxiang Wang, Chunyan Li, Haibo Peng, and Yalei Wang performed the experiments. Gang Chen and Jianye Zhang discussed the paper.

## Acknowledgments

This study was supported by The *National Natural Science Foundation of China* (Grant No. 31571369). We appreciate peers, colleagues and reviewers for their thoughtful comments and suggestions. Their insights helped us to push forward and deepen this study.

