## Supplemental text and Table S1-S12 for "A frameshift mutation is repaired through nonsense-mediated gene revising in *E. coli*"

### Supplementary text and tables for
### A frameshift mutation is repaired through nonsense-mediated gene revising in *E. coli*

*Xiaolong Wang[1*], Xuxiang Wang[1], Chunyan Li[1], Haibo Peng[1], Gang Chen[1], Jianye Zhang[1]*

*College of Life Sciences, Ocean University of China, Qingdao, 266003, P. R. China*

#### 1. Materials and Methods in details

##### *1.1 Frameshift preparation and revertant screening*

The G:C base pair located at +136 was deleted from the wild-type *bla* gene (*bla*+) by an overlapping extension polymerase chain reaction (OE-PCR). *E. coli* DH5α competent cells were transformed with pBR322-(bla-), propagated in tetracycline broth. Dilutions were plated in parallel on tetracycline plates and ampicillin plates to screen for revertants. The recovery rates were calculated by a standard method [1]. The revertants were propagated in ampicillin broth at 37°C with 200 rpm shaking and overnight seed culture (1:50). The growth rates of the revertants were evaluated by the doubling time in their exponential growth phase. Their plasmid DNA was extracted, and their *bla* genes were sequenced by the Sanger method.

##### *1.2 Construction and expression of a PTC-free frameshift*

Usually, a suppressor tRNA gene is introduced into cells to readthrough a nonsense mutation and incorporate an amino acid instead of terminating the translation. However, suppressor tRNA genes are not suitable for reading through the PTCs of *bla-* for two reasons: first, all three different PTCs (UGA, UAG, UAA) are present in *bla-*, we must introduce three different suppressor tRNA genes. However, there is no means to restrict their function to *bla* and not to other genes. If three suppressor tRNAs were introduced, all the true termination codons of other genes would also be readthrough, producing all sorts of odd peptides in the host; second, even if three different suppressor tRNAs were introduced, there is no guarantee that *bla-* would be expressed as expected, because the

[*] To whom correspondence should be addressed: Xiaolong Wang, Department of Biotechnology, Ocean University of China, No. 5 Yushan Road, Qingdao, 266003, Shandong, P. R. China, Tel: 0086-139-6969-3150,. .

nonsense mRNAs may be subject to nonsense-mediated mRNA decay or translational frameshifting pathways.

A PTC-free frameshift *bla*, denoted as *bla#*, was derived from *bla-* by replacing each nonsense codon with a sense codon according to the readthrough rules (Table 1). A stop codon, TAA, was added in the 3'-end. The *bla#* gene was chemically synthesized by *Sangon Biotech, Co. Ltd* (*Shanghai*), inserted into the expression vector pET28a and transformed into *E. coli* competent cells strain BL21. Since pET28a has a kanamycin resistance gene, the transformants were then plated on a kanamycin plate, propagated in kanamycin broth, and plated on ampicillin plates to screen for revertants. The expression of the *bla#* gene was induced by 0.1 mM IPTG. Their total protein samples were extracted, the product was purified by the nickel column chromatography and analyzed by sodium dodecyl sulfate-polyacrylamide gel electrophoresis (SDS-PAGE). The purified product was tested by an iodometry assay to measure its lactamase activity .

##### *1.3 Plasmid and genomic DNA extraction*

The revertant (DH5α/pBR322/*bla**) was transferred in ampicillin and tetracycline containing LB, and the frameshift (DH5α/pBR322/*bla-*) was inoculated in tetracycline containing LB. After inoculation, the cells were cultured at 37 °C and 200 rpm for 12 h. For each strain, 1.0 mL cultures were sent to *Sangon Biotech Co., Ltd* (*Shanghai*) to Sanger sequence the *bla* and *tet* genes. The sequencing primer was tet-f: TAA CGC AGT CAG GCA CCG t (65-83 base on plasmid pBR322). The genome DNA of the above strains were extracted using genome DNA extraction kit (Tiangen).

##### *1.4 Genome resequencing and structure/variation analysis*

Library preparation and genome sequencing were conducted by a commercial service provided by *Novogene Co. Ltd*. Paired-end reads were obtained on an Illumina HiSeq 250PE platform. The raw reads were trimmed by Trimmomatic (v0.39) to remove adapters and low-quality sequences for each sample. The clean reads were mapped onto the reference genome of *E. coli* K12 MG1655 (NC_000913.3) and the reference sequence of plasmid pBR322 by bwa (v0.7.17); the output alignments were indexed by Samtools (v0.1.18) [2], sorted by Picard (v2.23.4); then the reads around indels were realigned, and single nucleotide polymorphisms (SNPs) and indels were called by GATK (v4.1.2.0) [3]. Structural variations (SVs) were scanned by BreakDancer (v1.3.6) [4, 5].

##### *1.5 RNA extraction*

The wild type (DH5α/pBR322/*bla*+) and the revertants (DH5α/pBR322/*bla**) were inoculated in amp and tet containing LB, and the frameshift (DH5α/pBR322/*bla*-) was inoculated in tet containing LB. After inoculation, the cells were cultured at 37 °C and 200 rpm for 12 h. The wild type, the frameshift and the revertants were inoculated in fresh LB with appropriate antibiotics at the ratio of 1:100 and cultured at 37 °C and 200 rpm for 3 h to log phase. The total RNA samples of the above strains were extracted using RNA extraction kit (Tiangen).

##### *1.6 Transcriptome analysis*

Library preparation and RNA sequencing were conducted by a commercial service (*Novogene Co. Ltd*). The high-quality clean reads for each sample were mapped using bwa (v0.7.17) onto the reference sequence of the *E. coli* K12 MG1655 genome (NC_000913.3), and the plasmid pBR322 (J01749.1) using Circos (v0.69) [6], the coverage depths of the transcriptomic reads were displayed on a ring diagram next to those of the corresponding genomic reads.

The expression level for each gene was calculated and analyzed as followed:

(1) *Quantification of gene expression level*

Bowtie (v2-2.2.3) was used for aligning the clean reads to the reference genome [7]. HTSeq (v0.6.1) was used to count the numbers of reads mapped to each gene [8]. The expected number of Fragments PerKilobase of transcript sequence per million base pairs sequenced (FPKM) was calculated for each gene.

(2) *Identification of differential expression genes*

To identify differential expression genes (*DEGs*), read counts were adjusted by the edgeR program for each genotype [9]. Differential expressions of the read counts of two conditions were performed using the DEGSeq R package (1.20.0) [10]. The P values were adjusted by the Benjamini & Hochberg method. The significantly differential expression threshold was set as corrected P-value (q-value) $< 0.005$; fold change $\geq 2.0$ and $\geq 1.5$ are considered great and moderate changes.

(3) *GO and KEGG enrichment of the DEGs*

Gene Ontology (GO) enrichment of the DEGs was implemented by the GOseq R package [11], in which the potential gene length bias was corrected, and GO terms with corrected P-value < 0.05 were considered significantly enriched by DEGs. KOBAS 2.0 [12] was used to test the statistical enrichment of DEGs in the Kyoto encyclopedia of genes and genomes (KEGG) pathways (http://www.genome.jp/kegg/).

##### *1.7 Quantitative analysis of global proteome*

Quantitative analysis of global proteome was performed by *PTM-Biolabs* (*Hangzhou*)*, Co., Ltd*. Peptides were loaded onto a reverse-phase pre-column (*Acclaim PepMap 100*), separated on a reverse-phase analytical column and analyzed by Q Exactive™ hybrid quadrupole-Orbitrap mass spectrometer (*ThermoFisher Scientific*).

The MS/MS data were processed using the Mascot search engine (v.2.3.0). The tandem mass spectra were searched against the Uniprot database. GO annotation proteome was derived from the UniProt-GOA database (http://www.ebi.ac.uk/GOA/). Identified protein IDs were converted into UniProt ID and mapped to GO IDs. For the proteins unannotated in the UniProt-GOA database, InterProScan is used to determine their GO annotations.

#### 2. A review of previous studies that are supportive of the NMGR model

In the main text, the proposed NMGR model suggests that frame repair is triggered by PTCs, based on RNA-directed DNA repair, and nonsense mRNAs are edited before directing the repair of the coding DNA. NMGR integrates several key links, which have already been intensively studied, including mRNA surveillance, mRNA processing, mRNA editing, DNA recombination and repair. Like the connected pathways, frame repair should be widely existing and highly conserved among species. NMGR shows consistency with many previous studies. This supplementary text reviewed previous studies covering a wide range of prokaryotic and eukaryotic species to gain deeper knowledge.

##### *2.1 mRNA decay is linked to other nonsense mRNA processing pathways*

In eukaryotes, NMD uses the presence of an exon junction complex (EJC) downstream PTCs as a second signal to distinguish a PTC from a true stop codon [13]. In addition to NMD, the nonsense mRNAs may also be subject to some other pathways, including translational repression, transcriptional silencing, and alternative splicing, all of which are linked to NMD [14-16]. In eukaryotes, NMD regulating factors, including three interacting proteins, UPF1, UPF2 (also known as NMD2), and UPF3, are encoded by highly conserved genes originally identified in yeast [17].

A frameshifted gene is sometimes expressed through programmed translational frameshifting [18-20] or translational readthrough [21]. Translational frameshifting occurs in various organisms, from *E. coli* to mammals, whereby the ribosome is guided toward a triplet codon that is shifted one position to the upstream (+1) or downstream (−1) [22]. In yeast, telomere maintenance is globally controlled by the ribosomal frameshifting process and NMD [23]. Translational readthrough is a process wherein a nonsense mRNA is translated by translating the PTCs into specific amino acids [24]. Mutations in *upf* genes caused nonsense suppression and subsequent readthrough in yeast and 5-20 fold decreases in the rates of NMD [25], suggesting a direct link between NMD, nonsense suppression, and readthrough. The three Upf proteins all interact with Sup35, and Upf1 interact with a second release factor, Sup45 (eRF1), and binding of an SMG-1-Upf1-eRF1-eRF3 (SURF) complex to EJC triggers Upf1 phosphorylation and NMD [26]. Besides, the nonsense suppression caused by *upf* mutations is additive to those by *sup35* and *sup45* mutations [27], suggesting that nonsense suppression is a nonsense-mRNA processing pathway, an alternative to NMD. In a word, although nonsense mRNAs are usually subject to NMD, when NMD is inhibited, they can also be translated by ribosomal frameshifting or translational readthrough.

##### *2.2 RNA can direct DNA repair*

All types of RNAs are subject to either translation, degradation, or processing, and a variety of molecular/cellular mechanisms are involved in RNA processing [28]. In the last few decades, a link between DNA repair and RNA processing has been established. For example, several functional proteins respond to DNA damage, and base-excision repair enzymes, such as SMUG1, APE1, and PARP1, have been shown to participate in RNA surveillance and processing [29].

In eukaryotes, RNA is transcribed in the nucleus and transported to the cytoplasm [30]. Recently, there is growing evidence that RNAs can also be transported back to the nucleus to repair DNA double-strand breaks (DSBs), a process known as *RNA-directed DNA Repair* [31-33]. In yeast [33], bacteria, and human embryonic kidney cells [34], a synthetic DNA/RNA hybrid or an RNA-only oligonucleotide can serve as the template for repairing DSB in a homologous DNA. In *E. coli*, small RNA patches can direct DNA modification [35]. Here it is shown that mRNA could be edited and direct the repair of a frameshift mutation in its CDS. Four deaminase genes (*codA*, *cdd*, *add,* and *tadA*) and the mismatch repair pathways were upregulated in the frameshift or the revertant. Therefore, mismatch repair and RNA editing are probably both involved in the frame repair.

##### *2.3 RNA editing is linked to DNA repair*

RNA editing is widely existing in *E. coli* [36], plants [37], humans [38-40], and mammals [41]. There are several different forms of RNA editing, such as A-to-I editing [42], C-to-U editing [43], and the insertion/deletion of uridine [44], but the schematism are highly conserved among species, from bacteria to human. In particular, the nonsense *apoB* mRNA editing complex suppresses its NMD, suggesting that nonsense mRNAs can be subjected to RNA editing, which is linked to the NMD pathway [45].

Both DNA mismatch repair and RNA editing involve deamination, removing an amine group from a DNA/RNA molecule. Enzymes that catalyze this reaction are called DNA/RNA deaminases. Cytidine deamination was first recognized as a mechanism for RNA editing in wheat [46]. The tRNA-specific adenosine deaminases (ADATs) and the cytidine/adenosine deaminases (CDARs/ADARs) belong to a superfamily of RNA dependent deaminases [47]. Site-specific cytidine deamination introduces a stop codon (UAA) into the edited apolipoprotein mRNA [48]. It is well known that deamination and RNA editing play many important roles. In eukaryotes, studies have provided evidence that deamination is related to both DNA repair and RNA editing. For example, endonuclease V, an RNA editing enzyme highly conserved from *E. coli* to human [49], is involved in excision repair that removes deoxyinosine from DNA. Dysregulated RNA editing contributes to genome instability in cancer cells [50].

In *E. coli*, CodA catalyzes the hydrolytic deamination of cytosine to uracil, and the isoguanine, the oxidation product of adenine and isocytosine [51, 52]; Cdd scavenges cytidine and 2'-deoxycytidine for uridine monophosphate synthesis [53-55]; Add is an enzyme that converts adenine, adenosine, and deoxyadenosine to guanine [56, 57]; in eukaryotes, the cytosine deaminase (CDA) [58], the cytidine/deoxycytidine deaminase (APOBEC1) [36, 59], and the adenosine deaminase (ADAR1 and ADAR2) [60-62] are all responsible for RNA editing.

The tRNA-specific adenosine deaminase (TadA) was first identified in yeast [63] and is also a prokaryotic RNA editing enzyme that was reported first in *E. coli* [64]. TadA is also related to the eukaryotic RNA editing enzymes, ADAR1/ADAR2, which is widely existing in many eukaryotic species, such as humans [65], mouse [66], and fruit fly [67]. TadA, a prokaryotic deaminase for RNA editing first identified in *E. coli,* displays sequence similarity to the yeast tRNA deaminase subunit Tad2p [64]. Besides, cytidine deaminase (CodA) has been used as a molecular model for RNA editing and RNA substrate recognition [36]. Altogether, these studies suggest that RNA editing is highly conserved between prokaryotes and eukaryotes. Also, there is a direct link between the regulation of mismatch repair and RNA editing, supporting that both mismatch repair and RNA editing are involved in the frame repair.

### 3. Supplementary tables

Table S1. The number of changes in the DNA or RNA manipulating genes by comparing *Fs*, *Rs,* and *Rf* with *Wt* (see Table S2-S12 for the regulated genes in each pathway)

| | Total genes | Total comparisons | Total changes (>=1.2 fold) | | Great changes (>=2.0 fold) | | Moderate changes (1.2-2.0-fold) | |
|---|---|---|---|---|---|---|---|---|
| | | | up | down | up | down | up | down |
| DNA Replication | 18 | 54 | 16 | 2 | 3 | 0 | 13 | 2 |
| Transcription | 8 | 24 | 9 | 4 | 0 | 0 | 9 | 4 |
| RNA degradation | 16 | 48 | 25 | 2 | 5 | 0 | 20 | 2 |
| Nucleotide Excision Repair | 8 | 24 | 12 | 1 | 0 | 0 | 12 | 1 |
| Base Excision Repair | 14 | 42 | 4 | 5 | 0 | 0 | 4 | 5 |
| Mismatch Repair | 23 | 69 | 17 | 7 | 2 | 0 | 15 | 7 |
| Recombination | 28 | 84 | 29 | 7 | 6 | 1 | 23 | 6 |
| RNA editing | 10 | 30 | 7 | 3 | 2 | 0 | 5 | 3 |
| tRNA synthetases | 80 | 240 | 71 | 16 | 3 | 1 | 68 | 24 |
| RNA processing | 50 | 150 | 29 | 18 | 5 | 0 | 24 | 18 |
| **Translation** | **60** | **180** | **24** | **102** | **3** | **16** | **21** | **86** |
| **Total (exclude translation)** | **255** | **765** | **219** | **65** | **26** | **2** | **193** | **72** |
| Total (include translation) | 315 | 945 | 243 | 167 | 29 | 18 | 214 | 158 |

Table S2. The changes in the DNA replication pathways by comparing *Fs*, *Rs, Rf,* and *Wt.*

| Gene_id | FPKM | | | | Comparison | | | | | Gene | Description |
|---|---|---|---|---|---|---|---|---|---|---|---|
| | Wt | Fs | Rs | Rf | Fs-Wt | Rs-Wt | Rf-Wt | Rs-Fs | Rf-Fs | | |
| b1842 | 159.90 | 112.16 | 125.42 | 141.16 | -- | -- | -- | -- | -- | holE | DNA polymerase III subunit theta |
| b0215 | 170.54 | 190.38 | 125.74 | 165.32 | -- | -- | -- | ↓ | -- | dnaQ | DNA polymerase III subunit epsilon |
| b0184 | 66.08 | 116.03 | 87.85 | 93.12 | ↑ | ↑ | ↑ | -- | ↓ | dnaE | DNA polymerase III subunit alpha |
| b0470 | 121.51 | 156.36 | 152.15 | 158.94 | ↑ | ↑ | ↑ | -- | -- | dnaX | DNA polymerase III subunit tau |
| b1099 | 40.46 | 69.92 | 65.39 | 62.11 | ↑ | ↑ | ↑ | -- | -- | holB | DNA polymerase III subunit delta'. |
| b0640 | 37.00 | 83.50 | 82.68 | 56.94 | ↑↑ | ↑↑ | -- | -- | -- | holA | DNA polymerase III subunit delta |
| b4372 | 100.80 | 126.11 | 61.87 | 135.62 | -- | -- | -- | ↓ | -- | holD | DNA polymerase III subunit psi |
| b4259 | 64.79 | 69.41 | 55.30 | 64.80 | -- | -- | -- | -- | -- | holC | DNA polymerase III subunit chi |
| b3701 | 137.09 | 174.00 | 159.33 | 175.95 | -- | -- | ↑ | -- | -- | dnaN | beta sliding clamp |
| b4052 | 126.16 | 137.92 | 139.98 | 189.76 | -- | -- | ↑ | -- | ↑ | dnaB | replicative DNA helicase |
| b3066 | 185.31 | 189.31 | 173.12 | 187.35 | -- | -- | -- | -- | -- | dnaG | DNA primase |
| b4059 | 791.00 | 533.29 | 695.11 | 644.55 | ↓ | -- | -- | ↑ | -- | ssb | ssDNA-binding protein |
| b0214 | 136.98 | 127.92 | 97.39 | 128.54 | -- | -- | -- | -- | -- | rnhA | ribonuclease HI |
| b0183 | 52.35 | 97.98 | 74.57 | 83.24 | ↑ | -- | -- | -- | -- | rnhB | RNase HII |
| b3863 | 149.99 | 126.42 | 174.69 | 139.64 | -- | ↑ | -- | ↑ | -- | polA | DNA polymerase I |
| b2411 | 71.96 | 74.97 | 80.18 | 68.76 | -- | -- | -- | -- | -- | ligA | DNA ligase |
| b3647 | 23.85 | 15.74 | 21.94 | 10.57 | -- | -- | ↓ | -- | -- | ligB | DNA ligase |
| b0231 | 6.40 | 157.62 | 23.59 | 28.07 | ↑↑ | -- | -- | ↓↓ | ↓↓ | dinB | DNA polymerase IV |

Note: One arrow: moderate (1.2-2.0-fold) changes; Two arrows: great (>=2.0 fold) changes; "--" no change

Table S3. The changes in the RNA polymerase by comparing *Fs*, *Rs, Rf,* and *Wt.*

| **Gene_id** | **FPKM** | | | | **Comparison** | | | | | **Gene** | **Description** |
|---|---|---|---|---|---|---|---|---|---|---|---|
| | **Wt** | **Fs** | **Rs** | **Rf** | **Fs-Wt** | **Rs-Wt** | **Rf-Wt** | **Rs-Fs** | **Rf-Fs** | | |
| b1922 | 2.47 | 0.27 | 0.74 | 1.21 | -- | -- | -- | -- | -- | fliA | RNA polymerase sigma 28 (sigma F) factor |
| b2741 | 2,586.95 | 1,946.22 | 1,868.15 | 1,213.38 | ↓ | ↓ | ↓ | -- | ↓ | rpoS | RNA polymerase sigma S (sigma 38) factor |
| b3202 | 198.75 | 233.24 | 260.55 | 259.49 | -- | ↑ | ↑ | -- | -- | rpoN | RNA polymerase sigma 54 (sigma N) factor |
| b3295 | 4,782.59 | 3,990.28 | 2,754.60 | 4,265.86 | -- | ↓ | -- | ↓ | -- | rpoA | RNA polymerase alpha subunit |
| b3649 | 633.75 | 736.77 | 590.78 | 1,014.62 | -- | -- | ↑ | -- | ↑ | rpoZ | RNA polymerase omega subunit |
| b3987 | 658.35 | 844.08 | 848.12 | 930.53 | ↑ | ↑ | ↑ | -- | -- | rpoB | RNA polymerase beta subunit |
| b3988 | 635.27 | 825.14 | 881.82 | 906.31 | ↑ | ↑ | ↑ | -- | -- | rpoC | RNA polymerase beta prime subunit |
| b4293 | 45.59 | 72.95 | 76.15 | 77.16 | -- | -- | -- | -- | -- | fecI | RNA polymerase sigma-19 factor |

Note: One arrow: moderate (1.2-2.0-fold) changes; Two arrows: great (>=2.0 fold) changes; "--" no change

Table S4. The changes in the RNA degradation pathway by comparing *Fs*, *Rs, Rf,* and *Wt.*

| **Gene_id** | **FPKM** | | | | **Comparison** | | | | | **Gene** | **Description** |
|---|---|---|---|---|---|---|---|---|---|---|---|
| | **Wt** | **Fs** | **Rs** | **Rf** | **Fs-Wt** | **Rs-Wt** | **Rf-Wt** | **Rs-Fs** | **Rf-Fs** | | |
| b2830 | 484.58 | 333.42 | 360.63 | 384.86 | ↓ | -- | -- | -- | -- | rppH | RNA pyrophosphohydrolase |
| b1084 | 217.99 | 339.27 | 357.25 | 312.34 | ↑ | ↑ | ↑ | -- | -- | rne | ribonuclease E |
| b3780 | 381.39 | 408.22 | 463.69 | 538.39 | -- | ↑ | ↑ | -- | ↑ | rhlB | ATP-dependent RNA helicase RhlB |
| b2779 | 317.56 | 642.64 | 730.83 | 727.81 | ↑↑ | ↑↑ | ↑↑ | ↑ | -- | eno | enolase |
| b3164 | 886.73 | 925.08 | 954.49 | 967.27 | -- | ↑ | -- | -- | -- | pnp | polynucleotide phosphorylase |
| b0797 | 28.36 | 53.78 | 29.77 | 80.35 | ↑ | -- | ↑↑ | ↓ | ↑ | rhlE | ATP-dependent RNA helicase RhlE |
| b4179 | 198.74 | 197.25 | 172.61 | 232.92 | -- | -- | ↑ | -- | -- | rnr | RNase R |
| b3162 | 405.05 | 663.10 | 448.47 | 854.50 | ↑ | -- | ↑↑ | ↓ | ↑ | deaD | ATP-dependent RNA helicase DeaD |
| b3822 | 41.24 | 45.05 | 59.08 | 58.12 | -- | ↑ | ↑ | -- | -- | recQ | ATP-dependent DNA helicase RecQ |
| b3783 | 949.48 | 1,440.18 | 1,175.19 | 1,447.95 | ↑ | ↑ | ↑ | -- | -- | rho | transcription termination factor Rho |
| b3916 | 302.97 | 338.55 | 309.10 | 326.10 | -- | -- | -- | -- | -- | pfkA | 6-phosphofructokinase I |
| b0014 | 537.27 | 441.40 | 528.59 | 936.60 | -- | -- | ↑ | ↑ | ↑↑ | dnaK | chaperone protein DnaK |
| b4143 | 1,187.84 | 794.90 | 1,139.53 | 1,971.73 | ↓ | -- | ↑ | ↑ | ↑↑ | groL | chaperonin GroEL |
| b4172 | 2,403.91 | 2,004.11 | 1,774.57 | 2,712.58 | -- | -- | ↑ | -- | ↑ | hfq | RNA-binding protein Hfq |
| b2501 | 60.40 | 72.55 | 96.75 | 71.79 | -- | ↑ | -- | ↑ | -- | ppk | polyphosphate kinase |
| b0143 | 184.82 | 261.57 | 166.04 | 252.07 | ↑ | -- | ↑ | ↓ | -- | pcnB | poly(A) polymerase I |

Note: One arrow: moderate (1.2-2.0-fold) changes; Two arrows: great (>=2.0 fold) changes; "--" no change

Table S5. The changes in the Nucleotide Excision Repair pathway by comparing *Fs*, *Rs, Rf,* and *Wt.*

| **Gene_id** | **FPKM** | | | | **Comparison** | | | | | **Gene** | **Description** |
|---|---|---|---|---|---|---|---|---|---|---|---|
| | **Wt** | **Fs** | **Rs** | **Rf** | **Fs-Wt** | **Rs-Wt** | **Rf-Wt** | **Rs-Fs** | **Rf-Fs** | | |
| b1114 | 43.89 | 52.44 | 70.37 | 75.78 | -- | ↑ | ↑ | ↑ | ↑ | mfd | transcription-repair coupling factor |
| b4058 | 114.36 | 168.96 | 196.18 | 172.82 | ↑ | ↑ | ↑ | -- | -- | uvrA | excision nuclease subunit A |
| b0779 | 118.30 | 155.32 | 169.32 | 132.72 | ↑ | ↑ | -- | -- | -- | uvrB | excision nuclease subunit |
| b1913 | 39.04 | 70.35 | 63.80 | 55.65 | ↑ | ↑ | ↑ | -- | -- | uvrC | excision nuclease subunit |
| b3813 | 131.27 | 149.09 | 138.27 | 160.66 | -- | -- | ↑ | -- | -- | uvrD | ssDNA translocase and dsDNA helicase<br>DNA helicase II |
| b3863 | 149.99 | 126.42 | 174.69 | 139.64 | -- | ↑ | -- | ↑ | -- | polA | DNA polymerase I |
| b2411 | 71.96 | 74.97 | 80.18 | 68.76 | -- | -- | -- | -- | -- | ligA | DNA ligase |
| b3647 | 23.85 | 15.74 | 21.94 | 10.57 | -- | -- | ↓ | -- | -- | ligB | DNA ligase |

Note: One arrow: moderate (1.2-2.0-fold) changes; Two arrows: great (>=2.0 fold) changes; "--" no change

Table S6. The changes in the Base Excision Repair pathway by comparing *Fs*, *Rs, Rf,* and *Wt.*

| **Gene_id** | **FPKM** | | | | **Comparison** | | | | | **Gene** | **Description** |
|---|---|---|---|---|---|---|---|---|---|---|---|
| | **Wt** | **Fs** | **Rs** | **Rf** | **Fs-Wt** | **Rs-Wt** | **Rf-Wt** | **Rs-Fs** | **Rf-Fs** | | |
| b3635 | 36.39 | 24.23 | 28.57 | 41.55 | -- | -- | -- | -- | -- | mutM | DNA-formamidopyrimidine glycosylase |
| b0714 | 26.91 | 39.65 | 40.15 | 36.99 | -- | -- | -- | -- | -- | nei | endonuclease VIII |
| b1633 | 20.38 | 35.80 | 26.39 | 34.54 | -- | -- | -- | -- | -- | nth | endonuclease III |
| b2068 | 28.24 | 26.59 | 26.63 | 29.57 | -- | -- | -- | -- | -- | alkA | DNA-3-methyladenine glycosylase 2 |
| b3549 | 58.88 | 43.85 | 49.18 | 42.97 | -- | -- | -- | -- | -- | tag | 3-methyl-adenine DNA glycosylase I, constitutive |
| b3068 | 295.96 | 219.51 | 170.04 | 220.11 | ↓ | ↓ | ↓ | -- | -- | mug | stationary phase mismatch/uracil DNA glycosylase |
| b2580 | 112.98 | 88.47 | 70.92 | 116.75 | -- | ↓ | -- | -- | -- | ung | uracil-DNA glycosylase |
| b2961 | 52.78 | 68.97 | 60.90 | 84.12 | -- | -- | ↑ | -- | -- | mutY | adenine DNA glycosylase |
| b1749 | 137.97 | 136.94 | 104.86 | 132.67 | -- | -- | -- | -- | -- | xthA | exodeoxyribonuclease III |
| b2159 | 81.34 | 83.73 | 94.71 | 99.17 | -- | -- | -- | -- | -- | nfo | endonuclease IV |
| b3863 | 149.99 | 126.42 | 174.69 | 139.64 | -- | ↑ | -- | ↑ | -- | polA | DNA polymerase I |
| b2892 | 71.07 | 95.99 | 82.32 | 102.17 | ↑ | -- | ↑ | -- | -- | recJ | ssDNA-specific exonuclease RecJ |
| b2411 | 71.96 | 74.97 | 80.18 | 68.76 | -- | -- | -- | -- | -- | ligA | DNA ligase |
| b3647 | 23.85 | 15.74 | 21.94 | 10.57 | -- | -- | ↓ | -- | -- | ligB | DNA ligase |

Note: One arrow: moderate (1.2-2.0-fold) changes; Two arrows: great (>=2.0 fold) changes; "--" no change

Table S7. The changes in the Mismatch Repair pathway by comparing *Fs*, *Rs, Rf,* and *Wt.*

| Gene_id | FPKM | | | | Comparison | | | | | Gene | Description |
|---|---|---|---|---|---|---|---|---|---|---|---|
| | Wt | Fs | Rs | Rf | Fs-Wt | Rs-Wt | Rf-Wt | Rs-Fs | Rf-Fs | | |
| b2733 | 89.07 | 86.96 | 100.67 | 93.92 | -- | -- | -- | -- | -- | mutS | DNA mismatch repair protein MutS |
| b4170 | 55.45 | 75.62 | 63.57 | 77.37 | ↑ | -- | ↑ | -- | -- | mutL | DNA mismatch repair protein MutL |
| b2831 | 61.25 | 62.87 | 50.69 | 56.86 | -- | -- | -- | -- | -- | mutH | DNA mismatch repair protein MutH |
| b3813 | 131.27 | 149.09 | 138.27 | 160.66 | -- | -- | ↑ | -- | -- | uvrD | ssDNA translocase and dsDNA helicase - DNA helicase II |
| b2011 | 60.19 | 69.41 | 77.81 | 73.14 | -- | -- | -- | -- | -- | sbcB | exodeoxyribonuclease I |
| b2509 | 62.95 | 65.14 | 83.75 | 81.14 | -- | -- | -- | -- | -- | xseA | exodeoxyribonuclease VII subunit XseA |
| b0422 | 748.31 | 584.81 | 532.88 | 442.02 | -- | ↓ | ↓ | -- | -- | xseB | exodeoxyribonuclease VII subunit XseB |
| b1844 | 104.99 | 115.16 | 91.13 | 120.72 | -- | -- | -- | -- | -- | exoX | exonuclease X |
| b2892 | 71.07 | 95.99 | 82.32 | 102.17 | ↑ | -- | ↑ | -- | -- | recJ | ssDNA-specific exonuclease RecJ |
| b4059 | 791.00 | 533.29 | 695.11 | 644.55 | ↓ | -- | -- | ↑ | -- | ssb | ssDNA-binding protein |
| b0184 | 66.08 | 116.03 | 87.85 | 93.12 | ↑ | ↑ | ↑ | -- | ↓ | dnaE | DNA polymerase III (DpoIII) subunit alpha |
| b3701 | 137.09 | 174.00 | 159.33 | 175.95 | -- | -- | ↑ | -- | -- | dnaN | beta sliding clamp |
| b0470 | 121.51 | 156.36 | 152.15 | 158.94 | ↑ | ↑ | ↑ | -- | -- | dnaX | DNA polymerase III (DpoIII) subunit tau |
| b0640 | 37.00 | 83.50 | 82.68 | 56.94 | ↑↑ | ↑↑ | -- | -- | -- | holA | DNA polymerase III (DpoIII) subunit delta |
| b1099 | 40.46 | 69.92 | 65.39 | 62.11 | ↑ | ↑ | ↑ | -- | -- | holB | DNA polymerase III (DpoIII) subunit delta'. |
| b0215 | 170.54 | 190.38 | 125.74 | 165.32 | -- | -- | -- | ↓ | -- | dnaQ | DNA polymerase III (DpoIII) subunit epsilon |
| b1842 | 159.90 | 112.16 | 125.42 | 141.16 | -- | -- | -- | -- | -- | holE | DNA polymerase III (DpoIII) subunit theta |
| b4259 | 64.79 | 69.41 | 55.30 | 64.80 | -- | -- | -- | -- | -- | holC | DNA polymerase III (DpoIII) subunit chi |
| b4372 | 100.80 | 126.11 | 61.87 | 135.62 | -- | -- | -- | ↓ | -- | holD | DNA polymerase III (DpoIII) subunit psi |
| b2411 | 71.96 | 74.97 | 80.18 | 68.76 | -- | -- | -- | -- | -- | ligA | DNA ligase |
| b3647 | 23.85 | 15.74 | 21.94 | 10.57 | -- | -- | ↓ | -- | -- | ligB | DNA ligase |
| b3387 | 49.65 | 61.44 | 68.81 | 65.83 | -- | -- | -- | -- | -- | dam | DNA adenine methyltransferase |
| b3068 | 295.96 | 219.51 | 170.04 | 220.11 | ↓ | ↓ | ↓ | -- | -- | mug | stationary phase mismatch/uracil DNA glycosylase |

Note: One arrow: moderate (1.2-2.0-fold) changes; Two arrows: great (>=2.0 fold) changes; "--" no change

Table S8. The changes in the Homologous Recombination pathway by comparing *Fs*, *Rs, Rf,* and *Wt.*

| **Gene_id** | **FPKM** | | | | **Comparison** | | | | | **Gene** | **Description** |
|---|---|---|---|---|---|---|---|---|---|---|---|
| | **Wt** | **Fs** | **Rs** | **Rf** | **Fs-Wt** | **Rs-Wt** | **Rf-Wt** | **Rs-Fs** | **Rf-Fs** | | |
| b2892 | 71.07 | 95.99 | 82.32 | 102.17 | ↑ | -- | ↑ | -- | -- | recJ | ssDNA-specific exonuclease RecJ |
| b4059 | 791.00 | 533.29 | 695.11 | 644.55 | ↓ | -- | -- | ↑ | -- | ssb | ssDNA-binding protein |
| b3700 | 26.95 | 48.98 | 43.10 | 49.03 | ↑ | -- | ↑ | -- | -- | recF | DNA repair protein RecF |
| b2565 | 16.56 | 29.08 | 30.53 | 33.01 | -- | -- | -- | -- | -- | recO | DNA repair protein RecO |
| b0472 | 97.58 | 174.91 | 136.43 | 166.60 | ↑ | -- | ↑ | -- | -- | recR | DNA repair protein RecR |
| b2616 | 26.71 | 139.48 | 37.52 | 52.98 | ↑↑ | -- | ↑↑ | ↓↓ | ↓↓ | recN | recombination and repair protein |
| b2699 | 933.87 | 1,370.65 | 730.25 | 1,047.19 | ↑ | -- | ↑ | ↓ | ↓ | recA | DNA recombination/repair protein RecA |
| b3863 | 149.99 | 126.42 | 174.69 | 139.64 | -- | ↑ | -- | ↑ | -- | polA | DNA polymerase I (DpoI) |
| b1861 | 190.64 | 146.25 | 108.82 | 112.54 | -- | ↓ | ↓ | -- | -- | ruvA | Holliday junction branch migration complex subunit, RuvA |
| b1860 | 87.65 | 111.83 | 94.36 | 86.58 | -- | -- | -- | -- | -- | ruvB | Holliday junction branch migration complex subunit, RuvB |
| b1863 | 75.18 | 110.55 | 106.26 | 156.33 | -- | -- | ↑↑ | -- | -- | ruvC | crossover junction endodeoxyribonuclease RuvC |
| b3652 | 12.71 | 18.67 | 20.36 | 26.05 | -- | -- | ↑ | -- | -- | recG | ATP-dependent DNA helicase RecG |
| b2820 | 15.94 | 24.93 | 31.06 | 27.71 | ↑ | ↑↑ | ↑ | -- | -- | recB | exodeoxyribonuclease V subunit RecB |
| b2822 | 28.78 | 31.87 | 40.90 | 33.12 | -- | ↑ | -- | -- | -- | recC | exodeoxyribonuclease V subunit RecC |
| b2819 | 22.55 | 27.61 | 27.07 | 27.39 | -- | -- | -- | -- | -- | recD | exodeoxyribonuclease V subunit RecD |
| b0184 | 66.08 | 116.03 | 87.85 | 93.12 | ↑ | ↑ | ↑ | -- | ↓ | dnaE | DNA polymerase III (DpoIII) subunit alpha |
| b3701 | 137.09 | 174.00 | 159.33 | 175.95 | -- | -- | ↑ | -- | -- | dnaN | beta sliding clamp |
| b0470 | 121.51 | 156.36 | 152.15 | 158.94 | ↑ | ↑ | ↑ | -- | -- | dnaX | DNA polymerase III (DpoIII) subunit tau |
| b0640 | 37.00 | 83.50 | 82.68 | 56.94 | ↑↑ | ↑↑ | -- | -- | -- | holA | DNA polymerase III (DpoIII) subunit delta |
| b1099 | 40.46 | 69.92 | 65.39 | 62.11 | ↑ | ↑ | ↑ | -- | -- | holB | DNA polymerase III (DpoIII) subunit delta'. |
| b0215 | 170.54 | 190.38 | 125.74 | 165.32 | -- | -- | -- | ↓ | -- | dnaQ | DNA polymerase III (DpoIII) subunit epsilon |
| b1842 | 159.90 | 112.16 | 125.42 | 141.16 | -- | -- | -- | -- | -- | holE | DNA polymerase III (DpoIII) subunit theta |
| b4259 | 64.79 | 69.41 | 55.30 | 64.80 | -- | -- | -- | -- | -- | holC | DNA polymerase III (DpoIII) subunit chi |
| b4372 | 100.80 | 126.11 | 61.87 | 135.62 | -- | -- | -- | ↓ | -- | holD | DNA polymerase III (DpoIII) subunit psi |
| b3935 | 26.89 | 23.65 | 22.90 | 31.40 | -- | -- | -- | -- | -- | priA | primosome factor N' |

| | | | | | | | | | | | |
|---|---|---|---|---|---|---|---|---|---|---|---|
| b4201 | 1,837.23 | 1,116.01 | 627.55 | 1,311.36 | ↓ | ↓↓ | ↓ | ↓ | -- | priB | primosomal replication protein N |
| b0467 | 104.26 | 110.41 | 59.22 | 106.67 | -- | ↓ | -- | ↓ | -- | priC | primosomal replication protein N'‘. |
| b4362 | 102.60 | 121.04 | 91.27 | 103.98 | -- | -- | -- | -- | -- | dnaT | primosomal protein DnaT |

Note: One arrow: moderate (1.2-2.0-fold) changes; Two arrows: great (>=2.0 fold) changes; "--" no change

Table S9. The changes in the RNA editing pathway by comparing *Fs*, *Rs, Rf,* and *Wt.*

| | **FPKM** | | | | **Comparison** | | | | | | |
|---|---|---|---|---|---|---|---|---|---|---|---|
| **Gene_id** | **Wt** | **Fs** | **Rs** | **Rf** | **Fs-Wt** | **Rs-Wt** | **Rf-Wt** | **Rs-Fs** | **Rf-Fs** | **Gene** | **Description** |
| b0324 | - | 48.82 | - | - | ↑↑ | -- | -- | ↓↓ | ↓↓ | yahJ | putative Metallo-dependent hydrolase domain deaminase |
| b0337 | - | 48.76 | - | - | ↑↑ | -- | -- | ↓↓ | ↓↓ | codA | cytosine/isoguanine deaminase |
| b1623 | 224.72 | 282.87 | 377.11 | 191.6 | ↑ | ↑ | -- | ↑ | ↓ | add | adenosine deaminase |
| b1809 | 424.13 | 340.22 | 302.62 | 398.27 | -- | ↓ | -- | -- | -- | yoaB | putative reactive intermediate deaminase |
| b2065 | 472.63 | 410.44 | 419.79 | 522.81 | -- | -- | -- | -- | ↑ | dcd | deoxycytidine triphosphate deaminase dCTP deaminase |
| b2143 | 68.82 | 48.13 | 66.27 | 114.28 | -- | -- | ↑ | -- | ↑↑ | cdd | cytidine/deoxycytidine deaminase |
| b2559 | 50.74 | 56.47 | 41.01 | 68.16 | -- | -- | -- | -- | -- | tadA | tRNA-specific adenosine deaminase |
| b2883 | 8.21 | 6.99 | 5.62 | 5.42 | -- | -- | -- | -- | -- | guaD | guanine deaminase |
| b3113 | 31.42 | 36.74 | 89.69 | 79.58 | -- | ↑ | ↑ | ↑ | ↑ | tdcF | putative reactive intermediate deaminase |
| b3665 | 48.74 | 39.43 | 28.19 | 24.28 | -- | ↓ | ↓ | -- | ↓ | adeD | cryptic adenine deaminase |

Note: One arrow: moderate (1.2-2.0-fold) changes; Two arrows: great (>=2.0 fold) changes; "--" no change

Table S10. The changes in the tRNA synthetases by comparing *Fs*, *Rs, Rf,* and *Wt.*

| **Gene_id** | **FPKM** | | | | **Comparison** | | | | | **Gene** | **Description** |
|---|---|---|---|---|---|---|---|---|---|---|---|
| | **Wt** | **Fs** | **Rs** | **Rf** | **Fs-Wt** | **Rs-Wt** | **Rf-Wt** | **Rs-Fs** | **Rf-Fs** | | |
| b0026 | 306.93 | 419.6 | 301.51 | 369.16 | ↑ | -- | ↑ | ↓ | -- | ileS | isoleucyl-tRNA synthetase |
| b0058 | 23.95 | 50.26 | 35.6 | 42.8 | ↑ | -- | -- | -- | -- | rluA | dual specificity 23S rRNA pseudouridine(746) tRNA |
| b0144 | 40.8 | 49.76 | 43.83 | 42.89 | -- | -- | -- | -- | -- | gluQ | glutamyl-Q tRNA (Asp) synthetase |
| b0188 | 61.38 | 74.49 | 51.81 | 68.32 | -- | -- | -- | -- | -- | tilS | tRNA (Ile)-lysidine synthetase |
| b0191 | 39.88 | 30.16 | 20.05 | 26.38 | -- | -- | -- | -- | -- | arfB | alternative stalled-ribosome rescue factor peptidyl-tRNA |
| b0194 | 309.7 | 542.15 | 355.68 | 494.7 | ↑ | ↑ | ↑ | ↓ | -- | proS | prolyl-tRNA synthetase |
| b0195 | 18.06 | 46.02 | 34.68 | 53.44 | ↑ | -- | ↑ | -- | -- | tsaA | tRNA-Thr(GGU) m(6)t(6)A37 methyltransferase SAM-dependent |
| b0405 | 75.28 | 93.47 | 66.31 | 98.66 | -- | -- | -- | -- | -- | queA | S-adenosylmethionine:tRNA ribosyltransferase-isomerase |
| b0406 | 169.86 | 286.77 | 202.34 | 292.16 | ↑ | -- | ↑ | ↓ | -- | tgt | tRNA -guanine transglycosylase |
| b0423 | 84.07 | 113.79 | 89.61 | 148.74 | ↑ | -- | ↑ | -- | ↑ | thiI | tRNA s(4)U8 sulfurtransferase |
| b0481 | 52.16 | 72.79 | 41.96 | 63.94 | -- | -- | -- | -- | -- | ybaK | Cys-tRNA (Pro)/Cys-tRNA (Cys) deacylase |
| b0503 | 31.3 | 36.57 | 27.43 | 33.12 | -- | -- | -- | -- | -- | mnmH | tRNA 2-selenouridine synthase selenophosphate-dependent |
| b0526 | 130.04 | 127.6 | 110.89 | 182.04 | -- | -- | ↑ | -- | ↑ | cysS | cysteinyl-tRNA synthetase |
| b0642 | 315.69 | 438.23 | 358.17 | 338.07 | ↑ | ↑ | -- | -- | ↓ | leuS | leucyl-tRNA synthetase |
| b0661 | 252.85 | 270.25 | 188.68 | 309.7 | -- | -- | ↑ | ↓ | -- | miaB | tRNA -i(6)A37 methylthiotransferase |
| b0680 | 176.19 | 171.88 | 190.77 | 180.76 | -- | -- | -- | -- | -- | glnS | glutamyl-tRNA synthetase |
| b0885 | 38.54 | 43.71 | 46.61 | 46 | -- | -- | -- | -- | -- | aat | leucyl/phenylalanyl-tRNA -protein transferase |
| b0893 | 609.37 | 639.09 | 755.81 | 748.31 | -- | ↑ | ↑ | ↑ | -- | serS | seryl-tRNA synthetase |
| b0930 | 358.2 | 450.29 | 490 | 540.03 | ↑ | ↑ | ↑ | -- | ↑ | asnS | asparaginyl tRNA synthetase |
| b0969 | 121.08 | 120.15 | 103.85 | 147.42 | -- | -- | -- | -- | -- | tusE | mnm(5)-s(2)U34-tRNA 2-thiolation sulfurtransferase |
| b1133 | 262.28 | 336.36 | 324.27 | 421.98 | ↑ | ↑ | ↑ | -- | ↑ | mnmA | tRNA (Gln CLys CGlu) U34 2-thiouridylase |
| b1204 | 185.77 | 118.44 | 109.92 | 153.21 | ↓ | ↓ | -- | -- | -- | pth | peptidyl-tRNA hydrolase |
| b1210 | 195.52 | 211.12 | 186.07 | 208.77 | -- | -- | -- | -- | -- | hemA | glutamyl tRNA reductase |
| b1285 | 14.75 | 30.14 | 22.77 | 8.69 | ↑ | -- | -- | -- | ↓↓ | gmr | cyclic-di-GMP phosphodiesterase csgD regulator |
| b1344 | 123.89 | 96.88 | 84.74 | 124.26 | -- | ↓ | -- | -- | -- | ttcA | tRNA s(2)C32 thioltransferase iron sulfur cluster protein |

| | | | | | | | | | | | |
|---|---|---|---|---|---|---|---|---|---|---|---|
| b1637 | 373.4 | 321.29 | 364.12 | 383.75 | -- | -- | -- | -- | ↑ | tyrS | tyrosyl-tRNA synthetase |
| b1713 | 190.52 | 245.27 | 267.5 | 249.5 | ↑ | ↑ | ↑ | -- | -- | pheT | phenylalanine tRNA synthetase beta subunit |
| b1714 | 160.25 | 179.93 | 167.86 | 194.92 | -- | -- | -- | -- | -- | pheS | phenylalanine tRNA synthetase alpha subunit |
| b1715 | 122.33 | 69.79 | 51.03 | 112.37 | -- | -- | -- | -- | -- | pheM | phenylalanyl-tRNA synthetase operon leader peptide |
| b1719 | 1,539.10 | 1,195.20 | 981.46 | 1,262.97 | ↓ | ↓ | -- | -- | -- | thrS | threonyl-tRNA synthetase |
| b1787 | 120.15 | 151.89 | 104.45 | 132.51 | -- | -- | -- | -- | -- | yeaK | aminoacyl-tRNA editing domain protein |
| b1807 | 95.93 | 94.76 | 114.22 | 119.25 | -- | -- | -- | -- | -- | tsaB | tRNA (ANN) t(6)A37 threonylcarbamoyladenosine |
| b1866 | 330.92 | 329.02 | 377.33 | 426.73 | -- | ↑ | ↑ | ↑ | ↑ | aspS | aspartyl-tRNA synthetase |
| b1871 | 34.16 | 67.24 | 55.25 | 64.04 | ↑ | ↑ | ↑ | -- | -- | cmoB | tRNA (cmo5U34)-carboxymethyltransferase |
| b1876 | 225.4 | 207.83 | 187.97 | 232.99 | -- | -- | -- | -- | -- | argS | arginyl-tRNA synthetase |
| b2114 | 146.67 | 191.26 | 203.32 | 245.01 | ↑ | ↑ | ↑ | -- | ↑ | metG | methionyl-tRNA synthetase |
| b2140 | 31.47 | 14.29 | 12.86 | 11.77 | ↓ | ↓ | ↓ | -- | -- | dusC | tRNA -dihydrouridine synthase |
| b2268 | 53.2 | 50.89 | 40.22 | 49.96 | -- | -- | -- | -- | -- | rbn | RNase BN tRNA processing enzyme |
| b2318 | 77.1 | 47.56 | 54.54 | 53.4 | ↓ | -- | -- | -- | -- | truA | tRNA pseudouridine(38-40) synthase |
| b2400 | 389.02 | 420.84 | 369.91 | 460.92 | -- | -- | ↑ | -- | -- | gltX | glutamyl-tRNA synthetase |
| b2474 | 16.56 | 26.58 | 23.66 | 20.33 | -- | -- | -- | -- | -- | tmcA | elongator methionine tRNA (ac4C34) acetyltransferase |
| b2514 | 362.68 | 396.88 | 401.25 | 428.19 | -- | -- | ↑ | -- | -- | hisS | histidyl tRNA synthetase |
| b2517 | 168.35 | 232.82 | 217.34 | 250.76 | ↑ | ↑ | ↑ | -- | -- | rlmN | dual specificity 23S rRNA m(2)A2503 tRNA |
| b2530 | 836.44 | 685.94 | 342.26 | 513.2 | -- | ↓↓ | ↓ | ↓ | ↓ | iscS | cysteine desulfurase (tRNA sulfurtransferase) PLP-dependent |
| b2532 | 674.13 | 474.42 | 485.87 | 661.23 | ↓ | ↓ | -- | -- | ↑ | trmJ | tRNA mC32 CmU32 2'-O-methyltransferase SAM-dependent |
| b2559 | 50.74 | 56.47 | 41.01 | 68.16 | -- | -- | -- | -- | -- | tadA | tRNA -specific adenosine deaminase |
| b2575 | 56.06 | 45.75 | 43.57 | 60.96 | -- | -- | -- | -- | -- | trmN | tRNA 1(Val) (adenine(37)-N6)-methyltransferase |
| b2607 | 4,051.00 | 2,481.39 | 1,861.11 | 2,726.77 | ↓ | ↓ | ↓ | -- | -- | trmD | tRNA m(1)G37 methyltransferase SAM-dependent |
| b2697 | 375.34 | 449.79 | 493.98 | 510.45 | ↑ | ↑ | ↑ | -- | -- | alaS | alanyl-tRNA synthetase |
| b2745 | 110.44 | 126.37 | 101.12 | 135.68 | -- | -- | -- | -- | -- | truD | tRNA (Glu) pseudouridine(13) synthase |
| b2791 | 48.31 | 52.39 | 39.71 | 40.53 | -- | -- | -- | -- | -- | truC | tRNA (Ile1 CAsp) pseudouridine(65) synthase |
| b2812 | 101 | 62.02 | 58.89 | 58.34 | ↓ | ↓ | ↓ | -- | -- | tcdA | tRNA threonylcarbamoyladenosine dehydratase sulfur acceptor for CsdA |
| b2890 | 246.25 | 391.01 | 413.01 | 428.45 | ↑ | ↑ | ↑ | -- | -- | lysS | lysine tRNA synthetase constitutive |
| b2960 | 200.52 | 144.76 | 116.3 | 168.56 | ↓ | ↓ | -- | -- | -- | trmI | tRNA m(7)G46 methyltransferase SAM-dependent |

| | | | | | | | | | | | |
|---|---|---|---|---|---|---|---|---|---|---|---|
| b3056 | 91.15 | 115.33 | 120.91 | 132.71 | -- | ↑ | ↑ | -- | -- | cca | fused tRNA nucleotidyl transferase |
| b3064 | 171.1 | 154.86 | 127.18 | 198.44 | -- | -- | -- | -- | -- | tsaD | tRNA (ANN) t(6)A37 threonylcarbamoyladenosine |
| b3074 | 33.6 | 37.13 | 22.81 | 49.22 | -- | -- | -- | -- | -- | ygjH | putative tRNA binding protein putative tRNA corner chaperone |
| b3166 | 140.18 | 202.1 | 178.34 | 240.61 | ↑ | ↑ | ↑ | -- | -- | truB | tRNA pseudouridine synthase |
| b3260 | 441.56 | 438.69 | 419.33 | 531.59 | -- | -- | ↑ | -- | ↑ | dusB | tRNA -dihydrouridine synthase |
| b3282 | 86.46 | 114.07 | 97.12 | 71.51 | -- | -- | -- | -- | -- | tsaC | tRNA (ANN) t(6)A37 threonylcarbamoyladenosine |
| b3343 | 37 | 42.94 | 46 | 45.41 | -- | -- | -- | -- | -- | tusB | mnm(5)-s(2)U34-tRNA synthesis 2-thiolation protein |
| b3344 | 18.74 | 13.63 | 17.17 | 15.5 | -- | -- | -- | -- | -- | tusC | mnm(5)-s(2)U34-tRNA synthesis 2-thiolation protein |
| b3345 | 28.45 | 41.08 | 47.47 | 36.5 | -- | -- | -- | -- | -- | tusD | sulfurtransferase for 2-thiolation step |
| b3384 | 174.75 | 181.63 | 190.72 | 220.52 | -- | -- | ↑ | -- | -- | trpS | tryptophanyl-tRNA synthetase |
| b3442 | 1.66 | 0.67 | 0.3 | 0.44 | -- | -- | -- | -- | -- | yhhZ | putative Hcp1 family polymorphic toxin protein |
| b3470 | 386.2 | 305.59 | 224.77 | 494.75 | -- | ↓ | -- | -- | ↑ | tusA | mnm(5)-s(2)U34-tRNA 2-thiolation sulfurtransferase |
| b3559 | 222.19 | 319.84 | 373.8 | 395.32 | ↑ | ↑ | ↑ | ↑ | ↑ | glyS | glycine tRNA synthetase, beta subunit |
| b3560 | 306.48 | 375.35 | 390.89 | 473.39 | ↑ | ↑ | ↑ | -- | ↑ | glyQ | glycine tRNA synthetase, alpha subunit |
| b3590 | 43.41 | 47.34 | 58.69 | 82.32 | -- | -- | ↑↑ | -- | ↑ | selB | selenocysteinyl-tRNA-specific translation factor |
| b3606 | 59.19 | 30.64 | 27.95 | 51.87 | -- | ↓ | -- | -- | -- | trmL | tRNA Leu mC34 CmU34 2'-O-methyltransferase |
| b3651 | 19.04 | 36.7 | 35.59 | 43.72 | -- | -- | ↑ | -- | -- | trmH | tRNA mG18-2'-O-methyltransferase SAM-dependent |
| b3706 | 80.01 | 81.68 | 72.09 | 87.5 | -- | -- | -- | -- | -- | mnmE | tRNA U34 5-methylaminomethyl-2-thiouridine |
| b3741 | 158.32 | 184.44 | 142.91 | 237.2 | -- | -- | ↑ | -- | ↑ | mnmG | 5-methylaminomethyl-2-thiouridine modification at tRNA U34 |
| b3887 | 78.65 | 54.67 | 61.31 | 60.51 | -- | -- | -- | -- | -- | dtd | D-tyr-tRNA (Tyr) deacylase |
| b3965 | 144.03 | 112.67 | 80.71 | 149.03 | -- | ↓ | -- | ↓ | -- | trmA | tRNA m(5)U54 methyltransferase SAM-dependent |
| b4049 | 251.31 | 180.59 | 182.61 | 264.41 | ↓ | -- | -- | -- | ↑ | dusA | tRNA -dihydrouridine synthase A |
| b4129 | 130.2 | 306.45 | 207.26 | 524.48 | ↑↑ | ↑ | ↑↑ | ↓ | ↑ | lysU | lysine tRNA synthetase, heat-inducible |
| b4168 | 33.82 | 46.73 | 35.56 | 47.18 | -- | -- | -- | -- | -- | tsaE | tRNA (ANN) t(6)A37 threonylcarbamoyladenosine |
| b4171 | 460.47 | 627.65 | 516.78 | 787.5 | ↑ | -- | ↑ | -- | ↑ | miaA | delta(2)-isopentenylpyrophosphate tRNA -adenosine transferase |
| b4258 | 203.57 | 261.43 | 236.72 | 319.98 | ↑ | ↑ | ↑ | -- | ↑ | valS | valyl-tRNA synthetase |

Note: One arrow: moderate (1.2-2.0-fold) changes; Two arrows: great (>=2.0 fold) changes; "--" no change

Table S11. The changes in the RNA processing pathways by comparing *Fs*, *Rs, Rf,* and *Wt.*

| **Gene_id** | **FPKM** | | | | **Comparison** | | | | | **Gene** | **Description** |
|---|---|---|---|---|---|---|---|---|---|---|---|
| | **Wt** | **Fs** | **Rs** | **Rf** | **Fs-Wt** | **Rs-Wt** | **Rf-Wt** | **Rs-Fs** | **Rf-Fs** | | |
| b0059 | 28.59 | 56.31 | 41.81 | 45.35 | ↑↑ | ↑ | ↑ | ↓ | -- | rapA | RNA polymerase remodeling/recycling factor ATPase |
| b0069 | 26.16 | 30.34 | 23.58 | 29.17 | -- | -- | -- | -- | -- | sgrR | transcriptional DNA -binding transcriptional activator of sgrS sRNA |
| b0147 | 24.75 | 29.57 | 23.95 | 24.3 | -- | -- | -- | -- | -- | ligT | 2'-5' RNA ligase |
| b0183 | 52.35 | 97.98 | 74.57 | 83.24 | ↑ | -- | -- | -- | -- | rnhB | ribonuclease HII degrades RNA of DNA -RNA hybrids |
| b0214 | 136.98 | 127.92 | 97.39 | 128.54 | -- | -- | -- | -- | -- | rnhA | ribonuclease HI degrades RNA of DNA -RNA hybrids |
| b0225 | 135.57 | 116.08 | 70.28 | 98.75 | -- | ↓ | -- | -- | -- | yafQ | mRNA interferase toxin of toxin-antitoxin pair YafQ/DinJ |
| b0233 | 3.56 | 28.04 | 0.89 | 2.19 | ↑ | -- | -- | ↓ | ↓ | yafO | mRNA interferase toxin of the YafO-YafN toxin-antitoxin system |
| b0528 | 22.51 | 19.35 | 11.61 | 18.01 | -- | -- | -- | -- | -- | ybcJ | ribosome-associated protein putative RNA -binding protein |
| b0623 | 24,920.95 | 28,949.78 | 27,107.59 | 19,381.45 | -- | ↑ | ↓ | -- | ↓ | cspE | constitutive cold shock family, transcription antitermination protein |
| b0797 | 28.36 | 53.78 | 29.77 | 80.35 | ↑ | -- | ↑↑ | ↓ | ↑ | rhlE | ATP-dependent RNA helicase |
| b1045 | 48.88 | 54.4 | 64.51 | 37.55 | -- | -- | -- | -- | -- | ymdB | O-acetyl-ADP-ribose deacetylase RNase III inhibitor |
| b1084 | 217.99 | 339.27 | 357.25 | 312.34 | ↑ | ↑ | ↑ | -- | -- | rne | endoribonuclease RNA -binding protein RNA degradosome |
| b1267 | 138.69 | 111.26 | 99.57 | 134.5 | -- | -- | -- | -- | -- | yciO | putative RNA binding protein |
| b1285 | 14.75 | 30.14 | 22.77 | 8.69 | ↑ | -- | -- | -- | ↓↓ | gmr | cyclic-di-GMP phosphodiesterase csgD regulator |
| b1652 | 64.95 | 78.76 | 69.79 | 79.65 | -- | -- | -- | -- | -- | rnt | RNase T exoribonuclease T structured DNA 3' exonuclease |
| b1831 | 832 | 883.41 | 746.81 | 988.34 | -- | -- | ↑ | -- | -- | proQ | RNA chaperone putative ProP translation regulator |
| b1844 | 104.99 | 115.16 | 91.13 | 120.72 | -- | -- | -- | -- | -- | exoX | exodeoxyribonuclease 10 DNA exonuclease X |
| b2184 | 19.36 | 20.62 | 21.87 | 20.89 | -- | -- | -- | -- | -- | yejH | putative ATP-dependent DNA or RNA helicase |
| b2501 | 60.4 | 72.55 | 96.75 | 71.79 | -- | ↑ | -- | ↑ | -- | ppk | polyphosphate kinase component of the RNA degradosome |
| b2610 | 544.21 | 458.71 | 486.52 | 606.58 | -- | -- | -- | -- | ↑ | ffh | Signal Recognition Particle (SRP) component with 4.5S RNA (ffs) |
| b2620 | 319.86 | 232.86 | 261.15 | 376.9 | ↓ | -- | -- | -- | ↑ | smpB | tmRNA -binding trans-translation protein |
| b2630 | 217.76 | 156.62 | 99.51 | 155.05 | ↓ | ↓ | ↓ | ↓ | -- | rnlA | CP4-57 prophage RNase LS |
| b2666 | 17.87 | 66.66 | 36.66 | 38.38 | -- | -- | -- | -- | -- | yqaE | cyaR sRNA -regulated protein |
| b2754 | 30.53 | 11.02 | 10.54 | 6.12 | -- | -- | ↓ | -- | -- | ygbF | CRISPR adaptation ssRNA endonuclease |

| | | | | | | | | | | | |
|---|---|---|---|---|---|---|---|---|---|---|---|
| b2756 | 8.88 | 8.18 | 4.42 | 6.39 | -- | -- | -- | -- | -- | casE | CRISPR RNA precursor cleavage enzyme CRISP RNA (crRNA ) |
| b2757 | 7.63 | 7.27 | 4.71 | 3.62 | -- | -- | -- | -- | -- | casD | CRISP RNA (crRNA ) containing Cascade antiviral complex protein |
| b2758 | 7.32 | 7.01 | 5.66 | 4.63 | -- | -- | -- | -- | -- | casC | CRISP RNA (crRNA ) containing Cascade antiviral complex protein |
| b2759 | 6.62 | 4.06 | 2.19 | 5.78 | -- | -- | -- | -- | -- | casB | CRISP RNA (crRNA ) containing Cascade antiviral complex protein |
| b2760 | 6.24 | 4.94 | 1.29 | 1.85 | -- | ↓ | -- | -- | -- | casA | CRISP RNA (crRNA ) containing Cascade antiviral complex protein |
| b2782 | 22.2 | 45.57 | 25.24 | 18.68 | -- | -- | -- | -- | -- | mazF | mRNA interferase toxin antitoxin is MazE |
| b2830 | 484.58 | 333.42 | 360.63 | 384.86 | ↓ | -- | -- | -- | -- | rppH | RNA pyrophosphohydrolase |
| b3022 | 77.13 | 249.81 | 19.03 | 32.29 | ↑↑ | ↓ | ↓ | ↓↓ | ↓↓ | mqsR | GCU-specific mRNA interferase toxin of the MqsR-MqsA |
| b3083 | 20.29 | 16.82 | 4.49 | 9.41 | -- | -- | -- | -- | -- | higB | mRNA interferase toxin of the HigB-HigA toxin-antitoxin system |
| b3162 | 405.05 | 663.1 | 448.47 | 854.5 | ↑ | -- | ↑↑ | ↓ | ↑ | deaD | ATP-dependent RNA helicase |
| b3180 | 960.37 | 938.02 | 1,118.23 | 1,511.19 | -- | -- | ↑ | -- | ↑ | yhbY | RNA binding protein associated with pre-50S ribosomal subunits |
| b3205 | 89.31 | 126.95 | 158.06 | 141.74 | ↑ | ↑↑ | ↑ | -- | -- | yhbJ | adaptor protein for GlmZ/GlmY sRNA decay |
| b3229 | 1,199.97 | 998 | 1,017.64 | 1,345.83 | -- | -- | ↑ | -- | ↑ | sspA | stringent starvation protein phage P1 late gene activator |
| b3252 | 67.15 | 65.02 | 60.61 | 72.68 | -- | -- | -- | -- | -- | csrD | targeting factor for csrBC sRNA degradation |
| b3421 | 22.72 | 15.52 | 12.09 | 11.51 | -- | -- | ↓ | -- | -- | rtcB | RNA -splicing ligase |
| b3556 | 81,944.30 | 66,608.84 | 81,277.95 | 129,310.02 | -- | -- | ↑ | ↑ | ↑ | cspA | RNA chaperone and antiterminator cold-inducible |
| b3704 | 67.09 | 96.51 | 100.11 | 150.15 | -- | -- | ↑ | -- | -- | rnpA | protein C5 component of RNase P |
| b3780 | 381.39 | 408.22 | 463.69 | 538.39 | -- | ↑ | ↑ | -- | ↑ | rhlB | ATP-dependent RNA helicase |
| b3929 | 1,947.51 | 955.16 | 1,298.03 | 1,578.28 | ↓ | ↓ | -- | ↑ | ↑ | rraA | ribonuclease E (RNase E) inhibitor protein |
| b3995 | 471.31 | 316.44 | 388.5 | 360.43 | ↓ | -- | ↓ | -- | -- | rsd | stationary phase protein binds sigma 70 RNA polymerase subunit |
| b4128 | 4.19 | 1.98 | 0.59 | 0.59 | -- | -- | -- | -- | -- | ghoS | antitoxin of GhoTS toxin-antitoxin pair endonuclease |
| b4172 | 2,403.91 | 2,004.11 | 1,774.57 | 2,712.58 | -- | -- | ↑ | -- | ↑ | hfq | global sRNA chaperone HF-I host factor |
| b4179 | 198.74 | 197.25 | 172.61 | 232.92 | -- | -- | ↑ | -- | -- | rnr | exoribonuclease R RNase R |
| b4255 | 1,389.96 | 1,005.89 | 1,414.53 | 1,726.12 | ↓ | -- | ↑ | ↑ | ↑ | rraB | protein inhibitor of RNase E |
| b4331 | 83.51 | 58.71 | 42.97 | 85.46 | -- | ↓ | -- | -- | -- | kptA | RNA 2'-phosphotransferase |
| b4475 | 22.52 | 17.37 | 12.85 | 16.12 | -- | -- | -- | -- | -- | rtcA | RNA 3'-terminal phosphate cyclase |

Note: One arrow: moderate (1.2-2.0-fold) changes; Two arrows: great (>=2.0 fold) changes; "--" no change

Table S12. The changes in ribosomal proteins and elongation factors by comparing *Fs*, *Rs, Rf,* and *Wt.*

| Gene_id | FPKM | | | | Comparison | | | | | Gene | Description |
|---|---|---|---|---|---|---|---|---|---|---|---|
| | Wt | Fs | Rs | Rf | Fs-Wt | Rs-Wt | Rf-Wt | Rs-Fs | Rf-Fs | | |
| b0911 | 4,881.80 | 4,329.23 | 2,915.26 | 4,615.03 | -- | ↓ | -- | ↓ | -- | rpsA | 30S ribosomal subunit protein S1 |
| b0169 | 5,210.67 | 5,638.12 | 2,769.82 | 5,580.26 | -- | ↓ | -- | ↓ | -- | rpsB | 30S ribosomal subunit protein S2 |
| b3314 | 6,001.73 | 4,246.63 | 2,606.57 | 4,780.97 | ↓ | ↓ | -- | ↓ | -- | rpsC | 30S ribosomal subunit protein S3 |
| b3296 | 5,306.18 | 3,213.22 | 2,401.51 | 3,520.50 | ↓ | ↓ | ↓ | ↓ | -- | rpsD | 30S ribosomal subunit protein S4 |
| b3303 | 5,219.52 | 3,714.18 | 2,514.56 | 4,122.91 | ↓ | ↓ | -- | ↓ | -- | rpsE | 30S ribosomal subunit protein S5 |
| b4200 | 11,237.63 | 7,208.42 | 5,210.04 | 10,515.37 | ↓ | ↓ | -- | ↓ | ↑ | rpsF | 30S ribosomal subunit protein S6 |
| b3341 | 2,787.97 | 3,276.46 | 1,991.64 | 4,018.87 | ↑ | ↓ | ↑ | ↓ | ↑ | rpsG | 30S ribosomal subunit protein S7 |
| b3306 | 4,977.60 | 3,348.79 | 2,361.72 | 3,394.19 | ↓ | ↓ | ↓ | ↓ | -- | rpsH | 30S ribosomal subunit protein S8 |
| b3230 | 3,231.65 | 4,069.99 | 2,615.69 | 5,751.05 | ↑ | -- | ↑ | ↓ | ↑ | rpsI | 30S ribosomal subunit protein S9 |
| b3321 | 22,467.02 | 13,936.05 | 9,627.80 | 15,731.15 | ↓ | ↓↓ | ↓ | ↓ | -- | rpsJ | 30S ribosomal subunit protein S10 |
| b3297 | 3,177.73 | 1,869.21 | 1,464.44 | 1,955.16 | ↓ | ↓ | ↓ | -- | -- | rpsK | 30S ribosomal subunit protein S11 |
| b3342 | 5,673.02 | 6,743.22 | 4,115.87 | 8,204.61 | ↑ | -- | ↑ | ↓ | ↑ | rpsL | 30S ribosomal subunit protein S12 |
| b3298 | 4,492.67 | 3,143.27 | 2,213.90 | 3,229.96 | ↓ | ↓ | ↓ | ↓ | -- | rpsM | 30S ribosomal subunit protein S13 |
| b3307 | 7,059.01 | 4,755.04 | 3,306.83 | 4,364.90 | ↓ | ↓ | ↓ | ↓ | -- | rpsN | 30S ribosomal subunit protein S14 |
| b3165 | 12,132.52 | 8,313.65 | 7,112.08 | 13,508.37 | ↓ | ↓ | -- | -- | ↑ | rpsO | 30S ribosomal subunit protein S15 |
| b2609 | 5,413.63 | 3,419.57 | 3,122.36 | 4,268.17 | ↓ | ↓ | -- | -- | ↑ | rpsP | 30S ribosomal subunit protein S16 |
| b3311 | 2,210.32 | 2,002.85 | 965.72 | 2,191.57 | -- | ↓ | -- | ↓ | -- | rpsQ | 30S ribosomal subunit protein S17 |
| b4202 | 437.71 | 564.74 | 233.22 | 635.53 | -- | ↓ | ↑ | ↓↓ | -- | rpsR | 30S ribosomal subunit protein S18 |
| b3316 | 4,119.94 | 2,301.92 | 1,226.47 | 2,079.92 | ↓ | ↓↓ | ↓ | ↓ | -- | rpsS | 30S ribosomal subunit protein S19 |
| b0023 | 4,072.86 | 5,394.79 | 5,076.88 | 8,211.16 | ↑ | ↑ | ↑↑ | -- | ↑ | rpsT | 30S ribosomal subunit protein S20 |
| b3065 | 6,413.36 | 6,892.60 | 7,728.74 | 10,755.13 | -- | ↑ | ↑ | ↑ | ↑ | rpsU | 30S ribosomal subunit protein S21 |
| b3984 | 6,103.14 | 4,581.04 | 2,863.10 | 5,969.33 | ↓ | ↓ | -- | ↓ | ↑ | rplA | 50S ribosomal subunit protein L1 |
| b3317 | 6,337.07 | 4,698.36 | 2,991.13 | 5,119.89 | ↓ | ↓ | -- | ↓ | -- | rplB | 50S ribosomal subunit protein L2 |
| b3320 | 7,454.33 | 6,288.82 | 4,150.88 | 6,857.93 | -- | ↓ | -- | ↓ | -- | rplC | 50S ribosomal subunit protein L3 |
| b3319 | 5,038.70 | 3,948.32 | 2,498.27 | 4,141.10 | ↓ | ↓ | -- | ↓ | -- | rplD | 50S ribosomal subunit protein L4 |
| b3308 | 4,410.18 | 4,447.61 | 2,841.56 | 4,613.33 | -- | ↓ | -- | ↓ | -- | rplE | 50S ribosomal subunit protein L5 |

| | | | | | | | | | | | |
|---|---|---|---|---|---|---|---|---|---|---|---|
| b3305 | 4,552.18 | 3,194.18 | 2,148.00 | 3,513.49 | ↓ | ↓ | ↓ | ↓ | -- | rplF | 50S ribosomal subunit protein L6 |
| b3986 | 13,058.77 | 8,926.02 | 5,092.16 | 14,120.49 | ↓ | ↓↓ | -- | ↓ | ↑ | rplL | 50S ribosomal subunit protein L12 |
| b4203 | 2,492.01 | 3,476.38 | 2,210.96 | 5,343.81 | ↑ | -- | ↑↑ | ↓ | ↑ | rplI | 50S ribosomal subunit protein L9 |
| b3985 | 12,894.73 | 7,521.79 | 4,063.46 | 9,987.31 | ↓ | ↓↓ | ↓ | ↓ | ↑ | rplJ | 50S ribosomal subunit protein L10 |
| b3983 | 5,281.39 | 3,632.83 | 2,258.66 | 4,716.88 | ↓ | ↓↓ | -- | ↓ | ↑ | rplK | 50S ribosomal subunit protein L11 |
| b3231 | 9,684.41 | 6,314.89 | 4,156.59 | 8,192.87 | ↓ | ↓↓ | -- | ↓ | ↑ | rplM | 50S ribosomal subunit protein L13 |
| b3310 | 12,197.98 | 7,839.69 | 5,939.85 | 9,373.70 | ↓ | ↓ | ↓ | -- | ↑ | rplN | 50S ribosomal subunit protein L14 |
| b3301 | 4,621.93 | 3,765.91 | 2,659.19 | 4,165.60 | -- | ↓ | -- | ↓ | -- | rplO | 50S ribosomal subunit protein L15 |
| b3313 | 4,454.14 | 2,679.18 | 1,515.55 | 2,969.78 | ↓ | ↓↓ | ↓ | ↓ | -- | rplP | 50S ribosomal subunit protein L16 |
| b3294 | 3,115.94 | 3,442.61 | 2,294.70 | 4,595.33 | -- | -- | ↑ | ↓ | ↑ | rplQ | 50S ribosomal subunit protein L17 |
| b3304 | 3,463.77 | 2,540.58 | 1,638.32 | 2,575.63 | ↓ | ↓ | ↓ | ↓ | -- | rplR | 50S ribosomal subunit protein L18 |
| b2606 | 3,761.28 | 2,610.89 | 1,906.68 | 3,320.01 | ↓ | ↓ | -- | ↓ | ↑ | rplS | 50S ribosomal subunit protein L19 |
| b1716 | 6,859.38 | 7,758.93 | 4,163.07 | 8,864.45 | -- | ↓ | ↑ | ↓ | -- | rplT | 50S ribosomal subunit protein L20 |
| b3186 | 3,981.84 | 2,691.33 | 2,539.44 | 4,506.75 | ↓ | ↓ | ↑ | -- | ↑ | rplU | 50S ribosomal subunit protein L21 |
| b3315 | 4,123.22 | 3,258.41 | 1,780.89 | 3,293.10 | ↓ | ↓↓ | -- | ↓ | -- | rplV | 50S ribosomal subunit protein L22 |
| b3318 | 6,497.71 | 3,348.46 | 2,008.53 | 2,979.80 | ↓ | ↓↓ | ↓↓ | ↓ | -- | rplW | 50S ribosomal subunit protein L23 |
| b3309 | 4,845.90 | 3,252.68 | 2,153.53 | 3,016.29 | ↓ | ↓ | ↓ | ↓ | -- | rplX | 50S ribosomal subunit protein L24 |
| b2185 | 6,230.81 | 4,500.72 | 2,802.96 | 7,068.34 | ↓ | ↓ | ↑ | ↓ | ↑ | rplY | 50S ribosomal subunit protein L25 |
| b3185 | 4,542.69 | 3,340.60 | 2,608.09 | 5,462.19 | ↓ | ↓ | ↑ | -- | ↑ | rpmA | 50S ribosomal subunit protein L27 |
| b3637 | 12,798.35 | 9,145.76 | 6,655.57 | 12,464.02 | ↓ | ↓ | -- | ↓ | ↑ | rpmB | 50S ribosomal subunit protein L28 |
| b3312 | 597.48 | 219.80 | 81.89 | 127.14 | ↓↓ | ↓↓ | ↓↓ | ↓ | -- | rpmC | 50S ribosomal subunit protein L29 |
| b3302 | 147.00 | 77.42 | 20.61 | 39.72 | -- | ↓↓ | ↓ | ↓ | -- | rpmD | 50S ribosomal subunit protein L30 |
| b3936 | 19,592.77 | 12,202.68 | 9,576.80 | 18,996.32 | ↓ | ↓ | -- | -- | ↑ | rpmE | 50S ribosomal subunit protein L31 |
| b0296 | - | 2.23 | - | - | -- | -- | -- | -- | -- | ykgM | putative ribosomal protein |
| b1089 | 3,310.71 | 3,353.72 | 2,066.07 | 4,307.10 | -- | ↓ | ↑ | ↓ | ↑ | rpmF | 50S ribosomal subunit protein L32 |
| b3636 | 147.98 | 133.19 | 55.73 | 173.33 | -- | ↓ | -- | -- | -- | rpmG | 50S ribosomal subunit protein L33 |
| b3703 | 3,278.27 | 3,960.45 | 2,425.59 | 5,739.31 | ↑ | -- | ↑ | ↓ | ↑ | rpmH | 50S ribosomal subunit protein L34 |
| b1717 | 3,469.05 | 3,268.40 | 1,357.05 | 2,791.65 | -- | ↓↓ | -- | ↓↓ | -- | rpmI | 50S ribosomal subunit protein L35 |
| b3299 | - | - | - | - | -- | -- | -- | -- | -- | rpmJ | 50S ribosomal subunit protein L36 |
| b4506 | - | 1.39 | - | - | -- | -- | -- | -- | -- | ykgO | putative ribosomal protein |

| | | | | | | | | | | | |
|---|---|---|---|---|---|---|---|---|---|---|---|
| b3590 | 43.41 | 47.34 | 58.69 | 82.32 | -- | -- | ↑↑ | -- | ↑ | selB | selenocysteinyl-tRNA -specific translation factor |
| b3339 | 26.52 | 15.24 | 12.67 | 14.71 | -- | ↓ | -- | -- | -- | tufA | translation elongation factor EF-Tu 1 |
| b3980 | 109.39 | 80.67 | 45.62 | 91.67 | ↓ | ↓↓ | -- | ↓ | -- | tufB | translation elongation factor EF-Tu 2 |
| b0170 | 2,420.23 | 2,735.05 | 1,536.00 | 2,736.68 | -- | ↓ | ↑ | ↓ | -- | tsf | translation elongation factor EF-Ts |

Note: One arrow: moderate (1.2-2.0-fold) changes; Two arrows: great (>=2.0 fold) changes; "--" no change
